# A biotin ligation assay reveals a complex proxiome for HLA-A2 and implicates MIA3 in cell surface expression of MHC class I molecules

**DOI:** 10.1101/2025.09.08.674948

**Authors:** William Mitchell, Émeric Leclerc, Denis Faubert, Shijian Zhang, Jacques Thibodeau

## Abstract

Antigen presentation via MHC class I molecules (MHC-Is) is a turning point in the establishment of immune responses to endogenous threats, such as viruses. From their synthesis to their cell surface display, MHC-Is travel to many compartments, including the ER, Golgi, and endosomes. They come close to a plethora of molecules, some of which regulate directly or indirectly their folding and trafficking. While many of these proteins are well characterized, such as those found in the peptide loading complex, others remain to be discovered. The proxiome can be studied using proximity labeling assays, such as BioID, which relies on a biotin ligase fused to a bait of interest that biotinylates lysine residues on nearby proteins. These modified targets can then be purified and identified by mass spectrometry. By fusing BioID to the HLA-A2 cytoplasmic tail, we have applied this technique to the MHC-I antigen and identified 209 potential specific interactors in HEK293 cells, including PDZD8 (LYVAC) and MIA3 (Tango1). We knocked out MIA3 in HEK293 cells and measured an increase in the expression of MHC-I molecules, suggesting a role for this vesicle budding protein in the regulation of MHC-I trafficking. Notably, MHC-I crosslinking identified targets that connect reverse signalling to diverse metabolic processes. Altogether, our results underscore the promise of HLA-coupled biotin ligases as a powerful approach to dissect antigen presentation pathways.

## INTRODUCTION

Classical MHC class I molecules (MHC-Is) engage in a wide range of interactions to regulate their synthesis, trafficking, and antigen presentation functions. MHC-Is must capture peptides before they can interact with TCRs at the surface of CD8 T cells. On a structural basis, these peptides are needed to assure stable folding of the MHC-Is and maintenance of the quaternary structure involving the β2-microglobulin (β2M). Thus, in addition to ubiquitous chaperones involved in protein folding, MHC-Is depend on sophisticated mechanisms to achieve filling their peptide binding groove. These include TAP and ERp57, which are part of the peptide loading complex. This multiproteic ER complex detects, stabilizes, and induces the adoption of an open conformation of the peptide binding groove to permit the entry of an antigenic peptide. ^1,2^ Loaded MHC-Is, sent to the cell surface via the Golgi apparatus, undergo constant recycling through clathrin-dependent and independent endocytosis to either sorting or degradative compartments. ^3,4^ The many compartments involved in both biogenesis and transport of MHC-Is provide the backdrop to well-characterized interactions with regulators, such as Bap31, which plays a role in regulating MHC-Is’ exit from the ER, with the thioredoxin TMX1, implicated in the protein quality control process or with COPI which comes in close proximity to MHC-I-linked tapasin to regulate retrograde transport from the Golgi to the ER. ^5–7^ MHC-I’s important role in cross-presentation of exogenous antigens also implicates an additional set of regulators, such as Sec22b, which affects phagosome maturation. ^8^ A recent study has identified 2 more proteins interacting with MHC-I via pulldown: VAPA and ESYT1, while also confirming Bap31’s recruitment by the peptide loading complex. ^9^ Such findings suggest that many MCHI interactors remain to be discovered.

The identification of new potential protein-protein interactions (PPIs) is possible through proximity-based affinity purification techniques, such as BioID, in which a biotin ligase from *E.coli* is fused to a protein of interest to biotinylate the lysine residues of nearby proteins. Biotinylated targets of the enzyme can then be purified via pulldown using streptavidin, as biotin is an uncommon form of protein modification in eukaryotes. ^10,11^ Analysis by mass-spectrometry allows for the identification of proteins that were in close proximity to the molecule of interest in vivo. ^12^ BioID2, a later improvement on the original technique, replaces the *E. coli* enzyme by a smaller biotin-ligase found in the Gram-negative bacterium *Aquifex aeolicus*, thus improving biotinylation efficiency and selectivity of screening for protein-protein interactions. ^13^ A subsequent alternative to the technique came with the development of TurboID, a modified version of the original *E.coli* BirA enzyme, which allows for increased biotinylation efficiency. This significantly shortens the duration of incubation with biotin, thus allowing the characterization of more transient PPIs. ^14,15^

In this study, we sought to establish BioID2 as a valuable tool for probing the molecular environment of MHC molecules engaged in antigen presentation. We show that the biotin ligase, when fused to the intracytoplasmic domain of HLA-A2, was functional and allowed us to further characterize the proxiome of MHC-I molecules. Interestingly, the use of the more efficient TurboID biotin ligase showed that antibody crosslinking of MHC-I engages reverse signalling pathways associated with cellular metabolism in HEK293 cells.

## MATERIAL AND METHODS

### Antibodies and reagents

Pan-HLA class I antibody (W6/32) was produced in our lab. ^16^ The mouse monoclonal antibody SS 3A5-E2 to BioID2 and the rabbit monoclonal (EP892Y) to GM130 were purchased at Abcam (Toronto, ON). Goat anti-mouse IgG coupled to Alexa-fluor 647, streptavidin coupled to Alexa-fluor 488, Alexa-fluor 647-coupled CD71-specific antibody, APC-coupled W6/32 and IgG2a isotype control were purchased at Biolegend (San-Diego, CA). Goat anti-rabbit coupled to Alexa-fluor 488 was purchased from Thermo Fisher Scientific (Whitby, ON).

Protease inhibitor cocktail (Sigma, Oakville ON), phenylmethylsulfonyl fluoride (PMSF), dithiothreitol (DTT) (both purchased from Thermo Fisher Scientific), and benzonase (Sigma) were added to buffers used for immunoprecipitation of biotinylated proteins. Biotin was purchased from Sigma.

### Plasmids

The HLA-A2-BioID2 construct was generated by fusing the cytoplasmic domain of HLA-A2 to the *Aquifex aeolicus* biotin-ligase BioID2 with 2 repeats of GGGGS linker to separate the two. The HLA-DMα-BioID2 construct was generated in a similar manner, substituting HLA-A2 for HLA-DMα. ^17^ Both constructs were ordered from Thermo Fisher Scientific’s Geneart service in pcDNA3.1 backbones and were codon optimized for expression in human cells. The plasmid containing the hCas enzyme, 2 gRNAs specific to MIA3, as well as a GFP tag and neomycin resistance gene was purchased at Vectorbuilder (Chicago IL).

### Cell lines, transfections and CRISPR KO

Human embryonic kidney HEK293 cells were grown in DMEM containing 10% FBS (Wisent, St-Bruno, QC). K-562 lymphoblast cells ^18^ were grown in RPMI containing 10% FBS and 2mM L-Glutamine (all from Wisent). About 1×10^6^ cells were seeded in 10 cm culture plates and transfected for 48h with plasmid DNA using polyethyleneimine (Polyscience, Niles IL) in ExtremeMEM (Wisent).

To generate MIA3KO cells, the plasmid containing Cas9 and gRNAs was transfected into HEK293 cells as described above. GFP expression was monitored by flow cytometry to confirm efficient transfection. Cells were then cloned by limiting dilution under selection with G418 (Wisent). The clones were screened for gene expression by PCR or protein expression on western blots.

### PCR genotyping

Genomic DNA was obtained using a Monarch Genomic DNA purification kit (NEB, Whitby ON) according to the manufacturer’s instructions. Primers specific to the MIA3 locus (FWD: 5’AATCCAAGAGGGGCCTGGTA3’, REV: 5’CCCTGCATTGCCTTGTGATG3’) were purchased at Alpha DNA (Montréal QC). Kappa genotyping mix (Sigma, Oakville ON) was used to perform PCR per manufacturer’s instructions.

### Flow cytometry

For intracellular staining, cells were fixed in 4% paraformaldehyde and permeabilized in PBS with 0.1% saponin and 1% BSA. Cells were stained and analyzed on a BD FACSCanto II or BD FACS Symphony A1. Gating was limited to single cells in both FSC and SSC. Data was analyzed using FlowJo.

### Western blotting

Cells were harvested and lysed in buffer containing 1% protease inhibitor complex and 1% Triton X-100 (both from Sigma). Denaturing Laemmli buffer was added to post-nuclear supernatant. Samples were run on 10% acrylamide gels and transferred onto Amersham Protran 45um nitrocellulose membrane (Cytiva, Vancouver, BC). Membranes were blocked using skim milk and probed using relevant antibodies. Signal was revealed using SuperSignal West Pico PLUS Chemiluminescent Substrate (Thermo Fisher) on a GE Amersham AI600 imager (Cytivia, Vancouver BC).

### BioID

HEK293 cells were transiently transfected with the HLA-A2-BioID2- or HLA-A2-TurboID-encoding plasmid. HLA-DMα-BioID2 was also transfected separately to serve as a control. After 24h, biotin (Sigma) was added to culture media at a concentration of 70uM and cells were incubated for a further 24h (BioID2) or 60 minutes (TurboID). Expression of BioID2 was confirmed by flow cytometry prior to cell lysis. Initially, lysis was performed in cold RIPA buffer containing 1% NP-40 (Thermo Scientific) and 0.1% SDS via sonication (3x 10s at 30% amplitude) on ice. Supernatants were then cleaned by centrifugation at 13 000 rpm for 30 minutes at 4°C. Alternatively, PBS containing 1% Triton X-100 was used instead of RIPA to avoid lysis of the nucleus. Post-nuclear supernatants were obtained by centrifugation at 13 000 RPM for 15 minutes. For subsequent steps, 0.1% SDS was introduced to the PBS washing buffer described above to dissociate non-covalent protein complexes. Immunoprecipitation was then performed using 70ul of streptavidin-coupled sepharose bead slurry (Cytiva, Vancouver BC). Supernatant and beads were incubated for a minimum of 3 hours under agitation at 4°C. Beads were then cleaned three consecutive times in 1mL of the corresponding lysis buffer (either RIPA or PBS+Triton+SDS) and three more times in 50mM ammonium bicarbonate. Cleaned beads were then analyzed by mass spectrometry at the IRCM proteomics platform, as previously described. ^19^

### MS data analysis

The peak list files were generated with Proteome Discoverer (version 2.4) using the following parameters: minimum mass set to 500 Da, maximum mass set to 6000 Da, no grouping of MS/MS spectra, precursor charge set to auto, and minimum number of fragment ions set to 5. Protein database searching was performed with Mascot 2.6 (Matrix Science) against the Uniprot human protein database (April 4th, 2023) also containing the common contaminant proteins observed in BioID samples. The mass tolerances for precursor and fragment ions were set to 10 ppm and 0.05 Da, respectively. Trypsin was used as the enzyme allowing for up to 1 missed cleavage. Cysteine carbamidomethylation was specified as a fixed modification, and methionine oxidation as variable modification. Data interpretation was performed using Scaffold (version 4.8). Identified proteins were filtered through the CRAPome database to eliminate the most common contaminants found in BioID experiments using thresholds of 10 for average score, 40 for max score and 15% for positive CRAPome entries. ^20^ Further filtering was performed via the exclusion of non-specifically biotinylated proteins found in both HLA-A2-BioID2 transfected cells and mock transfected cells for each individual experiment. Hits present in both HLA-A2-BioID2 and YFP-BioID data were excluded. The YFP-BioID dataset was generated previously by our lab. ^19^ To generate the final list of proteins containing the most specific potential interactors, hits present in both HLA-A2-BioID2 and the HLA-DMα-BioID2 control were eliminated along with any protein not consistently found throughout a minimum of 7 of the 12 HLA-A2-BioID2 replicates. A STRING graph of the remaining proteins was generated using Cytoscape v.3.10.0. ^21^ Stringapp v. 2.0.1 was used for integration to the STRING database, which contains known PPIs and protein associations. ^22,23^ A confidence threshold of 0.3 was used to define meaningful interaction in STRING. Clustering was performed with Cytocluster v.2.0.1. using the DCU algorithm to predict closely associated groups of proteins based on their interaction score in STRING. ^24^ Clusters with a p-value < 0.05 were identified on the graph. Gene ontology enrichment analysis of the same remaining proteins was performed using ShinyGO v0.77 using the human genome as background. ^25^ GO terms containing between 20 and 2000 members were considered. A false discovery rate (FDR) cutoff of 0.005 was applied. Peptides known to bind the MHC-I groove were identified using the immune epitope database (IEDB). ^26^ For the comparison of gene enrichment between BioID data sets, PANTHER version 19.0 ^27,28^ was used over ShinyGO to gather data for all GO terms regardless of enrichment. Only GO terms with more than 80 members were considered for comparison to avoid bias towards very specific GO terms.

### Crosslinking of HLA-A2

Pan MHC-I antibody W6/32 was titrated to non-saturating concentrations by flow cytometry. Cells were incubated with the antibody or with an IgG2a isotype control on ice for 30 minutes. Cells were washed and incubated with a goat anti-mouse secondary antibody at 37°C for 1h in media supplemented with 50mM biotin. For TurboID experiments, cells were lysed and processed as described above.

### Confocal Microscopy and image analysis

Cells were plated on poly-L-lysine-coated coverslips and incubated at 37°C overnight in a 12-well plate. Coverslips were washed three times with PBS and cells were fixed in 4% paraformaldehyde (PFA). After washings with PBS, cells were permeabilized for 30 min using PBS containing 0.05% saponin and 1% BSA. Primary and secondary antibodies were incubated in permeabilization buffer. After a last series of washes in PBS, coverslips were mounted on microscope slides using fluoromount (Sigma). Image acquisition was performed on a Zeiss LSM 990 confocal microscope (Toronto ON) using the ZEN software. Object-based colocalization was performed using CellProfiler v4.2.8. ^29^. Both the Fiji software ^30^ and the Weka segmentation plugin ^31^ were used for image segmentation and object measurement within images. For GM130-stained slides, colocalization with MHC-I was determined by measuring MHC-I signal within GM130-bright regions. As a control, many random objects were generated within each MHC-I image. To approximate the distribution of intensities inside stained cells, only objects with a mean MHC-I intensity above 30 were kept and compared to the regions identified using GM130. A Mann-Whitney test was performed to assess significance.

## RESULTS

### Fusion of HLA-A2 to the biotin ligase of Aquifex aeolicus

In order to apply a BioID2 approach to the discovery of new PPIs involving MHC-Is, a fusion protein between HLA-A2 and the biotin-ligase from the gram-negative bacterium *Aquifex aeolicus* (hereafter called BioID2) was generated by recombining the DNA coding sequences into the pcDNA3.1 plasmid vector. BioID2 was fused at its N-terminus to the intracellular C-terminus of HLA-A2 (Figure. 1A). Two repeats of a serine/glycine linker (GGGGS) were inserted between the two proteins. This short nonpolar amino acid sequence confers a greater degree of movement freedom to the BioID2 enzyme, increasing its range of action and potentially limiting overcrowding around the cytoplasmic tail of HLA-A2. ^32^ (Figure 1A) As control, a similar construct was made between the DMα chain and BioID2 (HLA-DMα-BioID2). The isolated HLA-DMα-BioID2 chain will remain in the ER in the absence of HLA-DMβ, allowing us to discard irrelevant background proteins and some chaperones involved in ERAD, for example. ^33^ The mechanism by which BioID2 biotinylates proteins near HLA-A2’s intracytoplasmic domain and the method used to recover said target proteins for MS analysis is schematically detailed in figure 1B.

**Figure 1.**
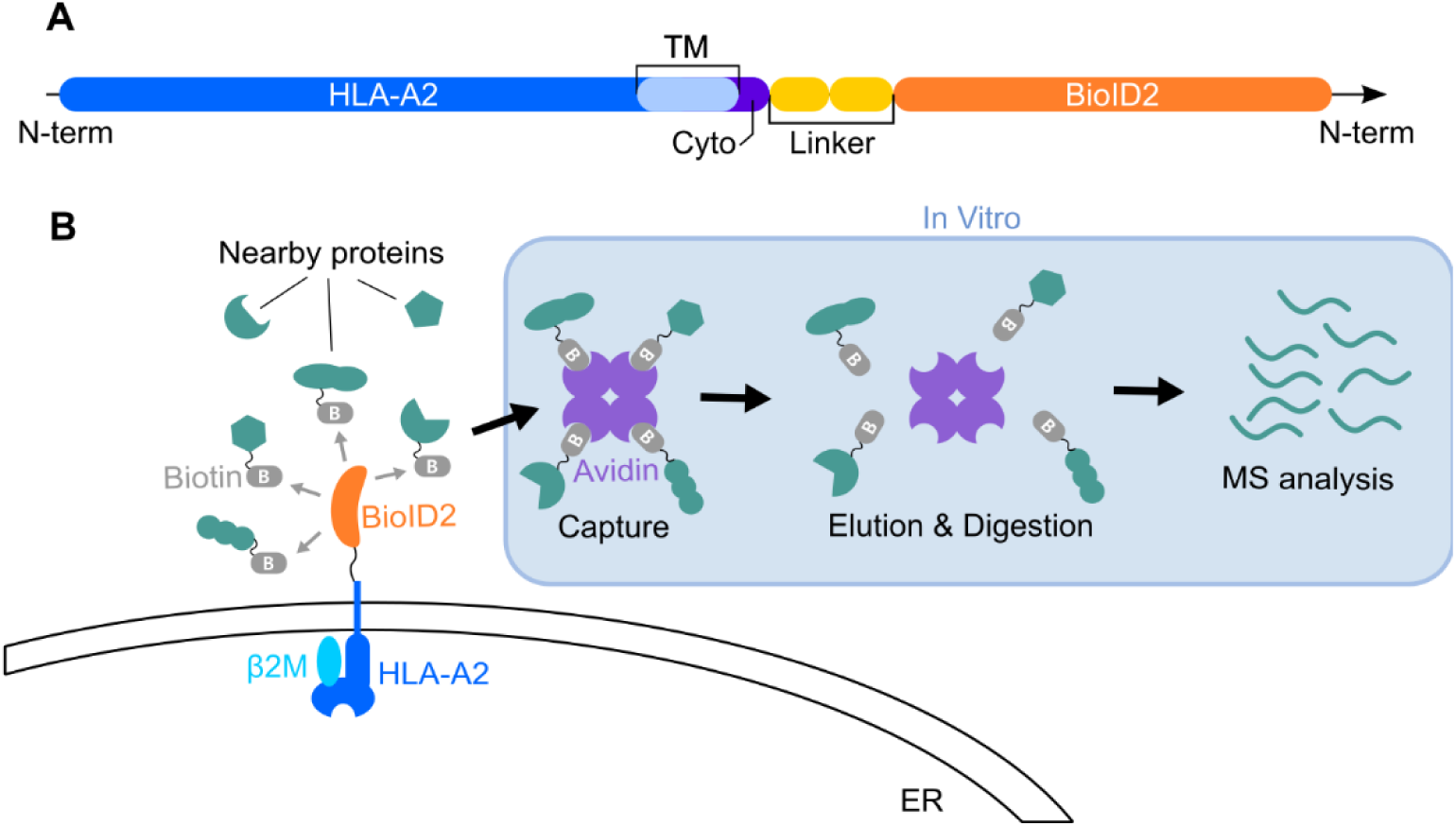
HLA-A2-adapted BioID2. **(A) Schematic representation of the HLA-A2-BioID2 fusion protein.** The HLA-A2 coding DNA sequence was fused at its C-terminus to the *A. aeolicus* biotin ligase. The HLA-A2 termination codon was changed for a flexible ser-gly linker. **(B) Principle of BioID2.** Along its journey through the ER, the secretory pathway, up the plasma membrane and the endocytic pathway, all molecules that come into a range of 10nm from the cytoplasmic biotin ligase will be ubiquitinated, given that an accessible lysine residue is exposed within this range.

### HLA-A2-BioID2 biotinylates nearby proteins

To assess the biotin ligase activity of HLA-A2-BioID2, the construct was transiently expressed in HEK293 cells. The size of the fusion protein and presence of the BioID2 enzyme were confirmed on immunoblots using an antibody targeted against BioID2. (Figure 2A) Levels of both the BioID2 enzyme and the accumulation of biotin-coupled intracellular proteins were measured by flow cytometry after a 16h incubation period of the cells with free biotin. Comparison between mock transfected cells and those expressing the fusion protein reveals the accumulation of intracellular biotin in transfected cells, in line with the successful labelling of proteins by HLA-A2 BioID2. (Figure 2B)

**Figure 2.**
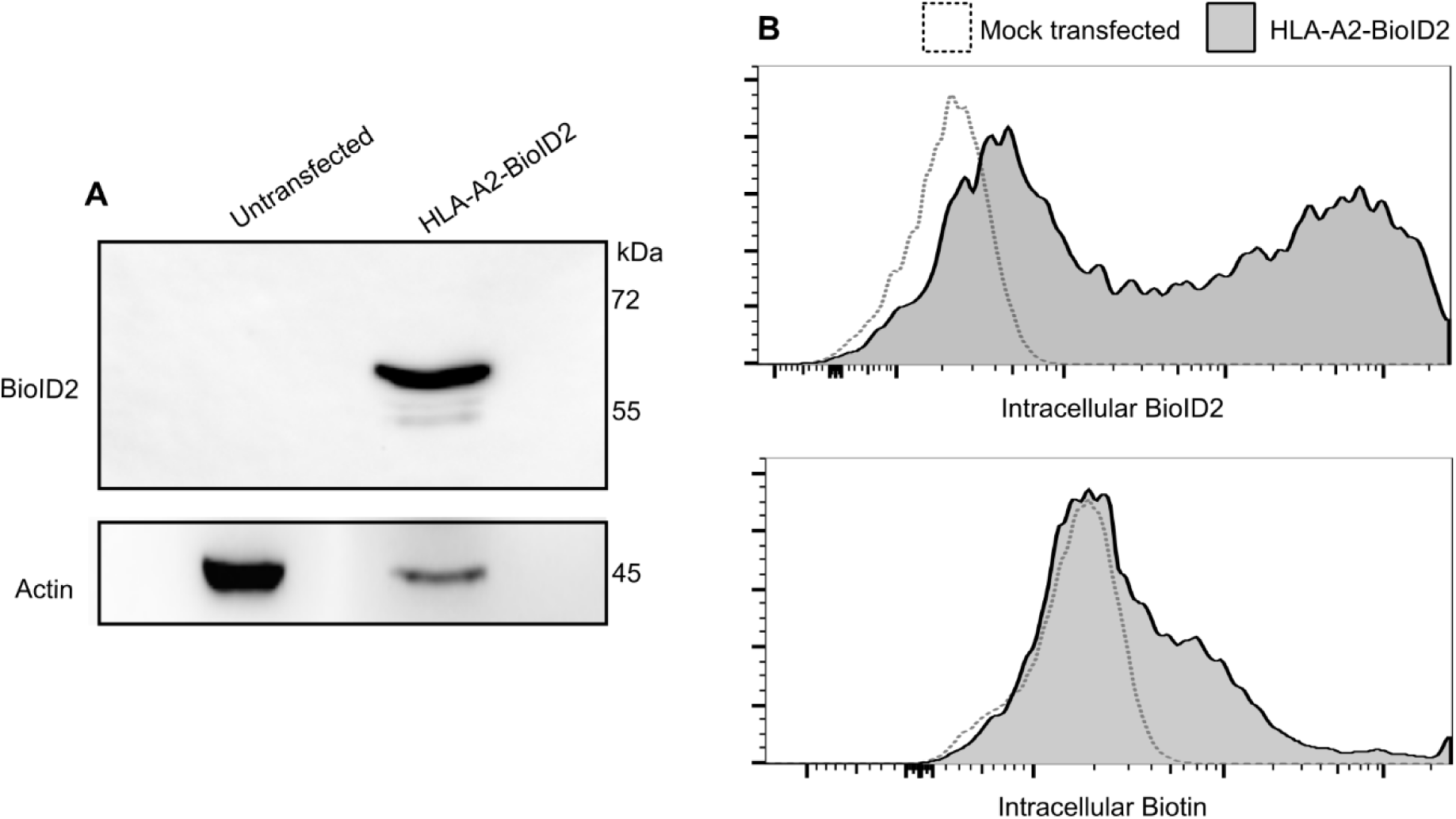
The HLA-A2-BioID2 fusion protein is functional and biotinylates intracellular targets. The HLA-A2-BioID2 construct was transfected into HEK293 cells. (A) Cells were lysed after 48h and the presence of the BioID2 and the molecular size of the construct were assessed by western blotting. Actin was probed as a loading control. (B) Flow cytometry analysis of cells 16h after incubation with biotin shows a marked increase in intracellular biotin and confirmed the presence of the BioID2 enzyme.

### HLA-A2-BioID2 is delivered at the plasma membrane

A previous study has shown that GFP could be fused to the C-terminus of HLA-A2 without disrupting its capacity to transit normally through the endocytic pathway. ^34^ Still, we ascertained that HLA-A2-BioID2 was folded properly and could reach the cell surface. The K-562 lymphoblastoid cell line, which lacks expression of MHC-I due to negative transcriptional regulation by the PRC2 complex ^18^ was transfected with HLA-A2-BioID2 and analyzed by flow cytometry for class I expression at the cell surface. The results show that both the control HLA-A2 and the HLA-A2-BioID2 accumulate at the plasma membrane. (Figure 3)

**Figure 3.**
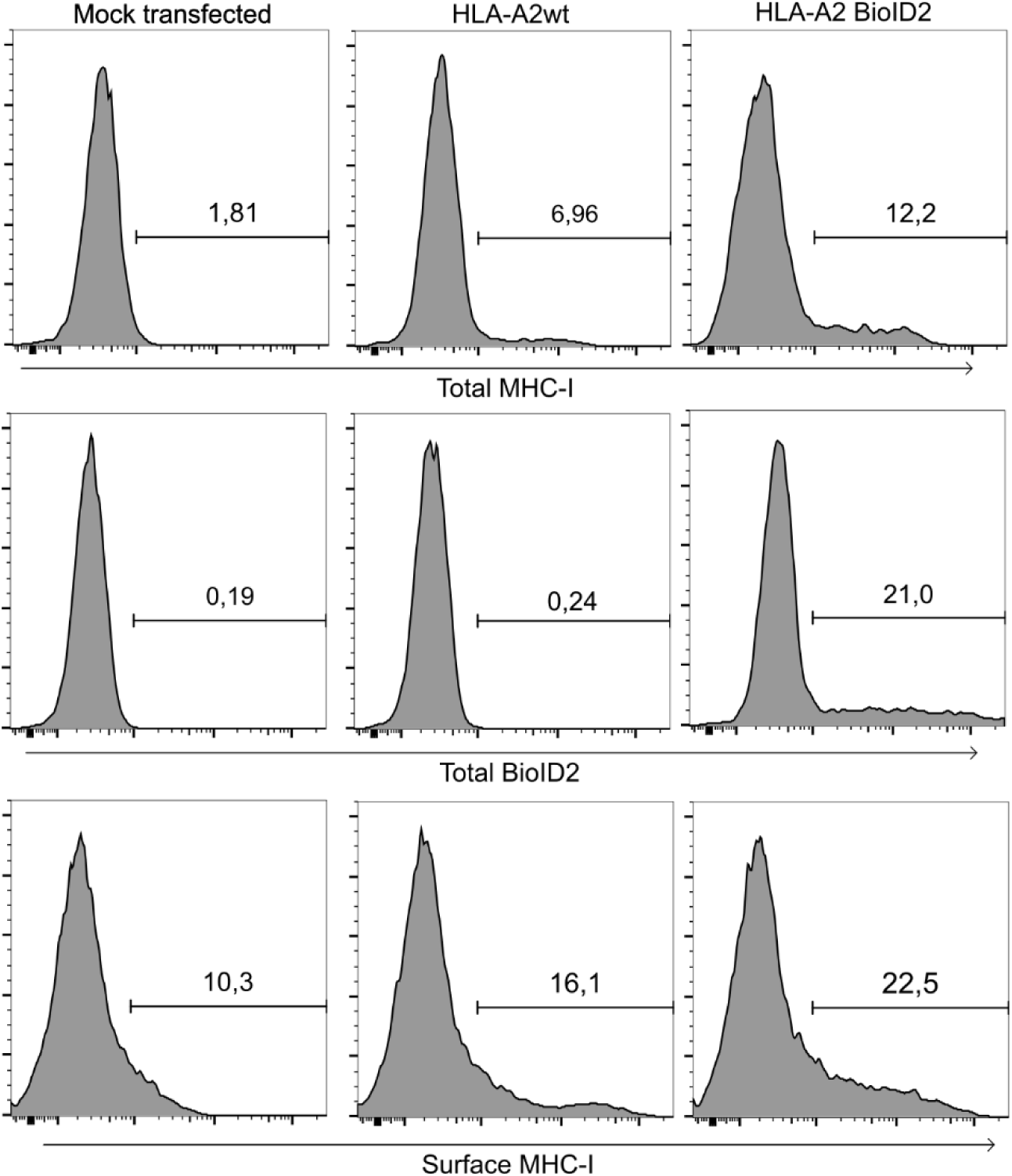
The HLA-A2 BioID2 fusion is expressed similarly to WT HLA-A2. RSV-HLA-A2 and pcDNA-HLA-A2-BioID2 were transiently transfected in the MHC-I negative K-562 cell line. After 48h, cells were surface-stained or fixed, permeabilized and stained (total). Expression of HLA-A2 and BioID2 was monitored by flow cytometry.

### A pulldown of biotinylated proteins coupled to mass spectrometry analysis allows for the identification of proteins near HLA-A2-BioID2

HLA-A2-BioID2 transfection, biotinylated protein pulldown and mass spectrometry analysis were performed in triplicate for each independent experiment. A total of 7017 targets were identified by mass spectrometry across 12 HLA-A2-BioID2 samples. (Figure 4A) Within the reproducible hits, which are defined as those present in at least 7 of the 12 HLA-A2-BioID samples, we found proteins already known to interact with HLA-A2, including 62 of the 264 HLA-A interactors described in the BioGRID database. (Figure 4B). ^35^ Calnexin, an important player of protein folding, was identified in 11 of 12 replicates, therefore showing HLA-A2-BioID2’s capacity to biotinylate membrane proteins. (Table S1) ^1^ Different fragments of HLA-B were also found in all 12 replicates as MHC-Is tend to form clusters on the cell surface. ^36^ Other BioGRID PPIs may not have been identified in our data as, for example, the BioID2 enzyme was located on the intracytoplasmic side and could not access proteins on the lumenal side of the many membranous organelles or the plasma membrane known to anchor MHC-Is. Conversely, many of the proteins in our dataset are not in the BioGRID list of interactors.

**Figure 4.**
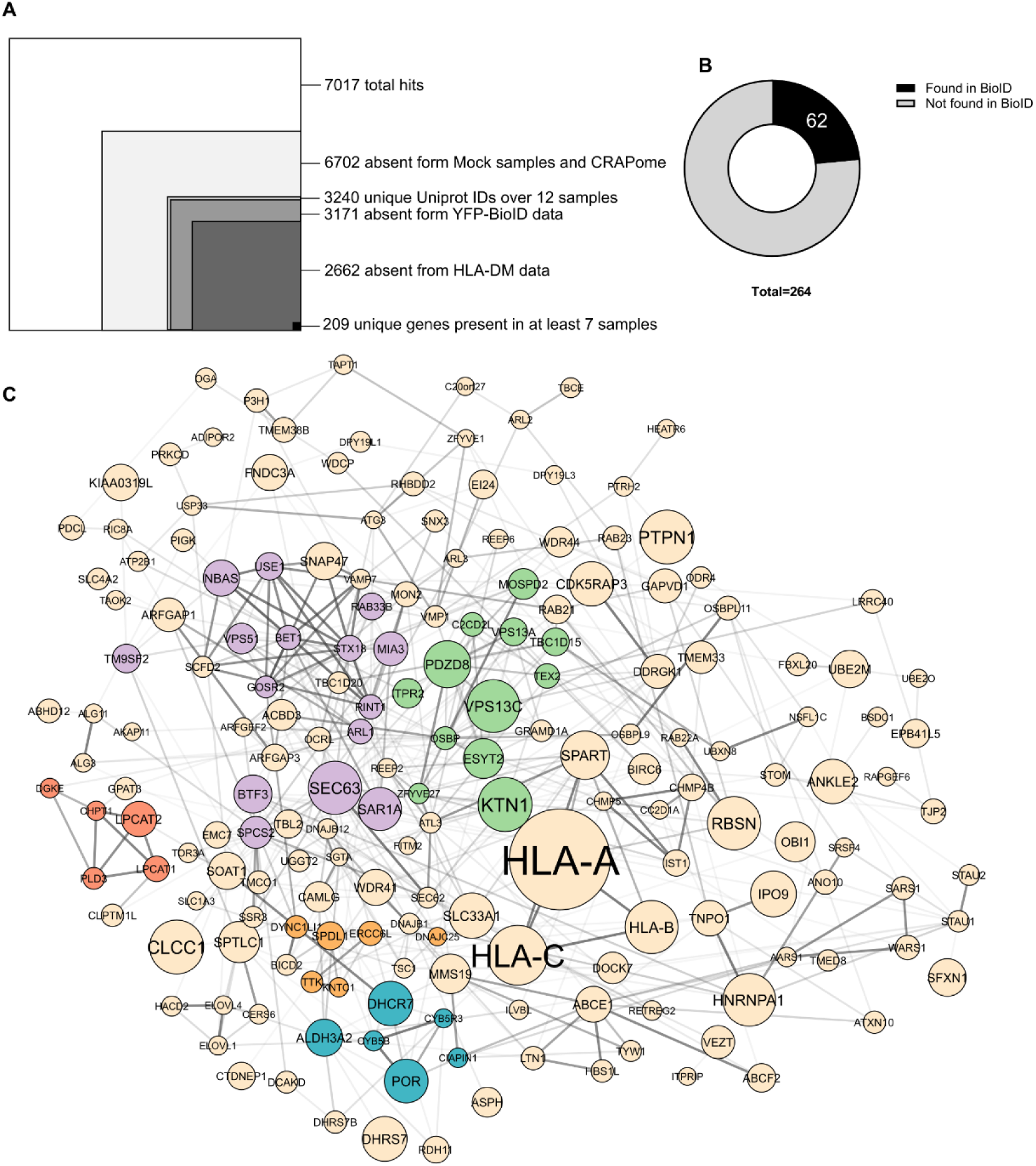
Filtering strategy applied to proteins identified by pulldown-MS for HLA-BioID2. **(A) Schematic representation of HLA-A2 MS data filtering**. All identified proteins from 12 HLA-A2-BioID2 samples were first filtered through the CRAPome database. All proteins present in mock transfected samples were discarded, as they most likely result from non-specific binding. The YFP-BioID dataset was then used to screen for proteins with greater scaffold quantitative scores in HLA-A2-BioID than in YFP-BioID. The HLA-DMα-BioID control served to eliminate most MHC-specific ER proteins not specific to HLA-A2. Hits consistently discovered in 7 of 12 replicates are considered reproducible. **(B) Proportion of BioGRID HLA-A PPIs found in BioID samples.** HLA-A2 BioID data was cross-referenced with the BioGRID database of known protein interactions. All human proteins known to interact with HLA-A were considered. Hits present in both HLA-A2-BioID and HLA-DMα-BioID2 were considered, as many interactions are known to happen in the ER. **(C) STRING Graph of proteins retained in 4A.** The graph was generated using Cytoscape v.3.10.0. Stringapp v. 2.0.1 was used for integration to the STRING database. Clustering was performed using Cytocluster v.2.0.1. which revealed 5 statistically significant clusters (p-value < 0.05). Further analysis revealed a common function of vesicular transport in the orange cluster and protein folding in the purple cluster. Line thickness and opacity represent the strength of protein associations within the STRING database, while node size represents the mean abundance of a protein in our HLA-A2-BioID2 data.

To improve the specificity of our analysis, we eliminated all proteins found using mock-transfected cells or listed in the CRAPome database. In addition, we discarded all proteins identified in previous experiments using a control YFP-BioID2 fusion protein transiently expressed in HEK293 cells. ^19^ Finally, data was further filtered as described in figure 4A to retain only specific, non-redundant and reproducibly discovered targets of HLA-BioID2 that were absent in HLA-DMα-BioID2 data. This process narrowed the original list of 7017 hits down to 209 candidate interacting proteins with unique UniProt IDs. (Tables S1,S2) These proteins are plotted in graphical form in figure 4C along with STRING database interactions. As HLA-A2 and other MHC-Is are biotinylated and were pulled-down by streptavidin, there is the possibility that some of the identified peptides are part of the immunopeptidome and have simply been released from the antigen-binding groove. Further analysis via the IEDB database of the individual peptides found by mass spectrometry suggested that only four proteins, including MHC-I isoforms themselves and CIAPIN1, were identified in part by peptides capable of binding the MHC-I groove. All groove peptides from the IEDB found in our raw data are detailed in table S3. Gene ontology (GO) enrichment analysis of the final 209 potential interactors was then performed using Shiny GO with the human genome as background. ^25^ Figure 5 details enriched GO terms with very significant FDR (<0.005) in HLA-A2 BioID experiments for all three main GO categories. Analysis of cell component GO terms reveals significant enrichment in terms related to the ER, Golgi, and endosomes, while biological process GO terms highlight an enrichment in proteins related to vesicle/membrane fusion and protein transport, which are indirectly related to the antigen presentation process. This further supports the notion that HLA-A2-BioID2 undergoes normal trafficking and that the proxiome likely encompasses functionally important proteins.

**Figure 5.**
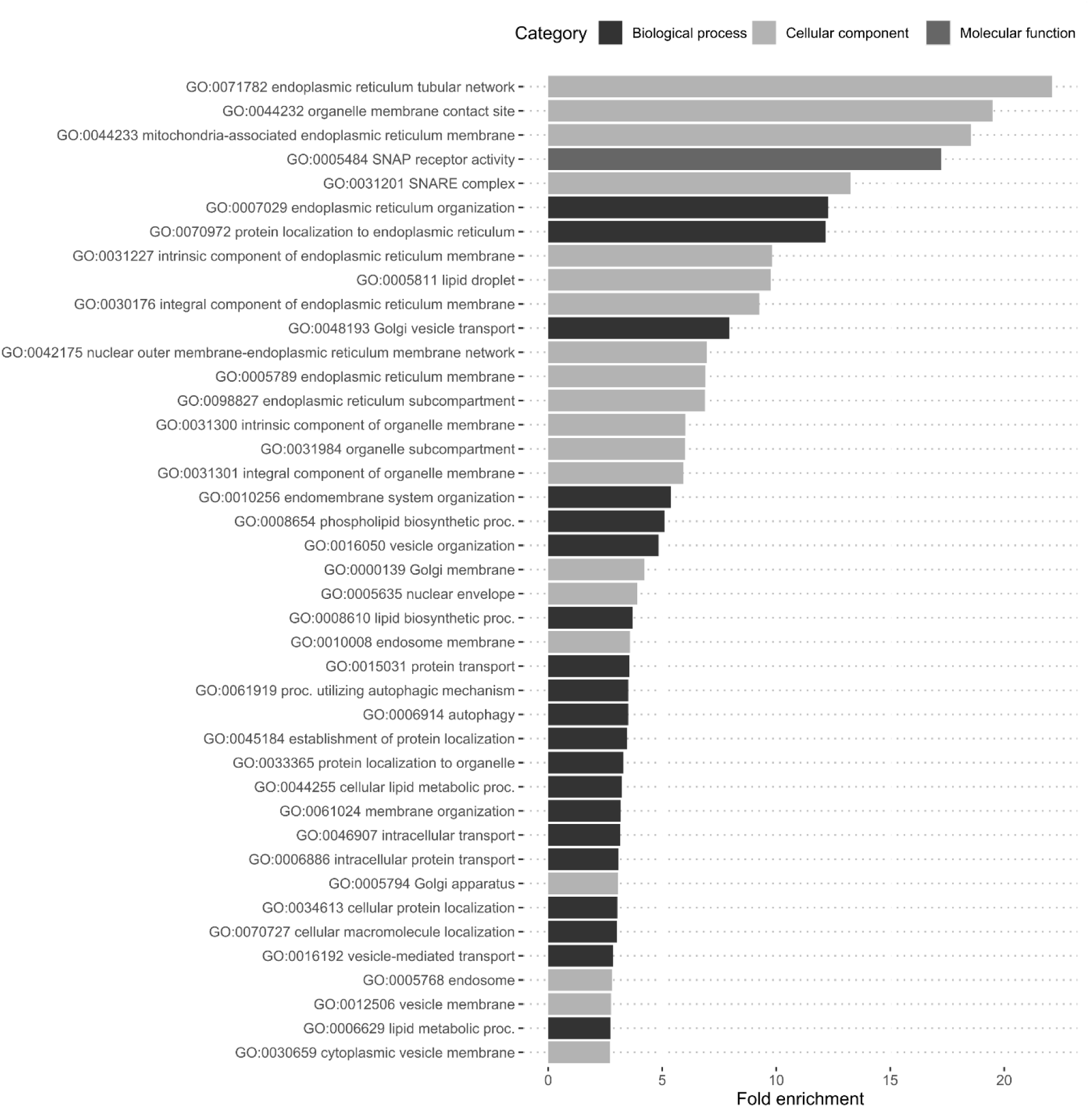
Gene ontology enrichment analysis of the PPIs. Gene ontology enrichment analysis of HLA-A2-BioID2 retained proteins was performed using ShinyGO v0.77. All significantly enriched (FDR>0.005) terms are ranked by fold enrichment. Bar colours indicate the main GO category for each term as per the legend.

### MIA3 knockout cells show higher expression of MHC-I and altered Golgi morphology

MIA3 (TANGO1), which was found in 10/12 BioID replicates, is implicated in endosome budding and is known to play a role in the formation of COPII vesicles, which can transport MHC-I. ^37–39^ To assess the possible role MIA3 plays in MHC-I antigen presentation, HEK293 MIA3KO cells were generated via CRISPR-Cas9 and clones were obtained by limiting dilutions. A MIA3 deletion in both genes of clone P3F12 was confirmed by PCR on purified genomic DNA. (Figure S1) Surface and total MHC-I expression was then measured by flow cytometry, revealing a global increase in MIA3KO cells as compared to WT cells. (Figure 6A) To determine if this effect was specific to MHC-Is, transferrin receptor (TFR; CD71) expression was also monitored, as it is a membrane protein that traffics in a manner similar to HLA-A2. ^40,41^ The results showed no variation of TFR for both surface and total expression between WT and MIA3KO cells. (Figure 6B) To gain further insights into this increased HLA class I expression, an internalisation assay was performed. However, no striking difference was observed between WT and MIA3KO cells. (Figure S2A) Accordingly, no significant difference was found by confocal microscopy in the colocalization of MHC-I and the EEA1 early endosome marker between WT and MIA3KO cells. (Figure S2BC) A similar experiment was conducted by co-staining the Golgi marker GM130 and MHC-Is in both cell lines. (Figure 7A) While colocalization of MHC-I within the identified Golgi objects (defined as GM130-bright regions) was immediately noticeable and similar, statistical analysis shows a significantly higher MHC-I presence in the Golgi of MIA3KO cells (Mann-Whitney p<0.0001). (Figure 7B,C) We theorized that this effect might be due to the overall higher expression of MHC-I in MIA3KO cells, but normalization using the randomized controls did not counterbalance the observed effect. Differences in Golgi morphology were however noted with MIA3KO cells where this compartment was significantly smaller after thresholding and particle analysis of a total of 340 GM130-bright objects across both cell lines. (Figure 7D) It is possible that similar quantities of MHC-I are present in the Golgi of both cell lines, but a smaller Golgi in MIA3KO makes for more a more concentrated and intense MHC-I signal.

**Figure 6.**
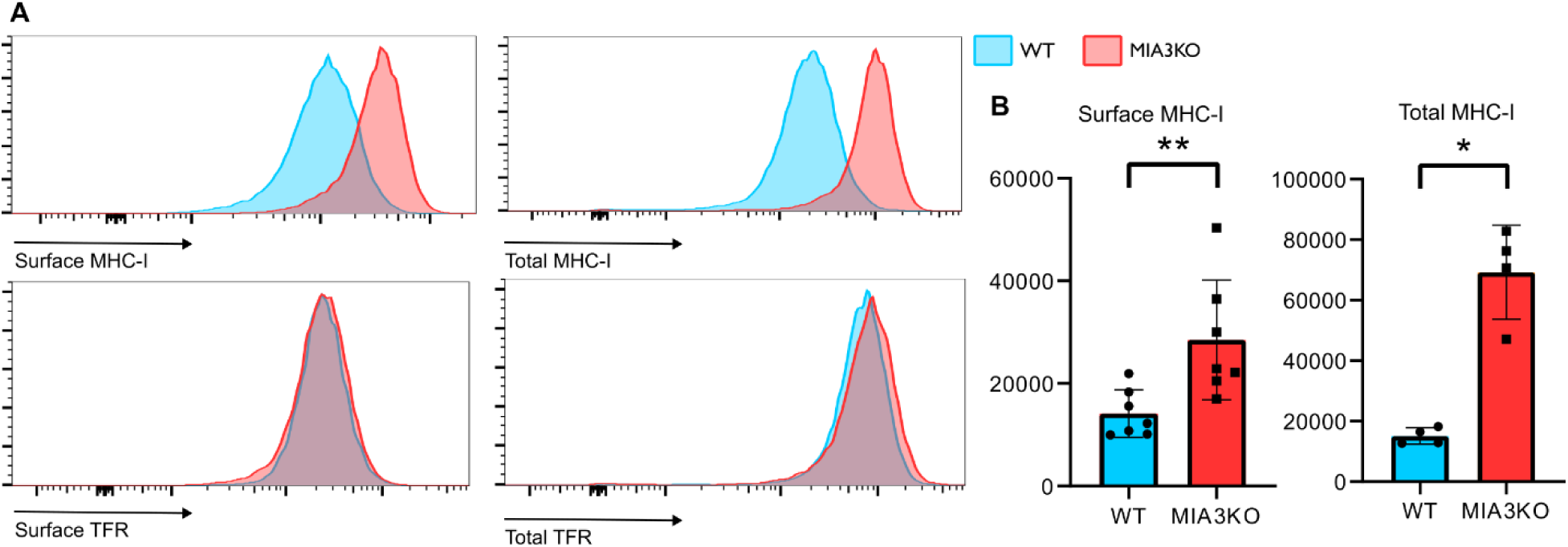
MIA3 negatively affects MHC-I expression. (A) Flow cytometry analysis of surface and total MHC-I as well as TFR in WT vs MIA3KO cells reveals a heightened presence of MHC-I in MIA3KO cells with no effect on the control membrane protein TFR. (B) Geometric means of fluorescence intensities (MFI) of 7 flow cytometry experiments comparing surface and 4 comparing total expression of MHC-I for WT and MIA3KO cells are represented in histograms. Statistical significance was determined via a Man-Whitney test (*p<0.05, **p<0.01) showing significant increases in MHC-I for both conditions.

**Figure 7:**
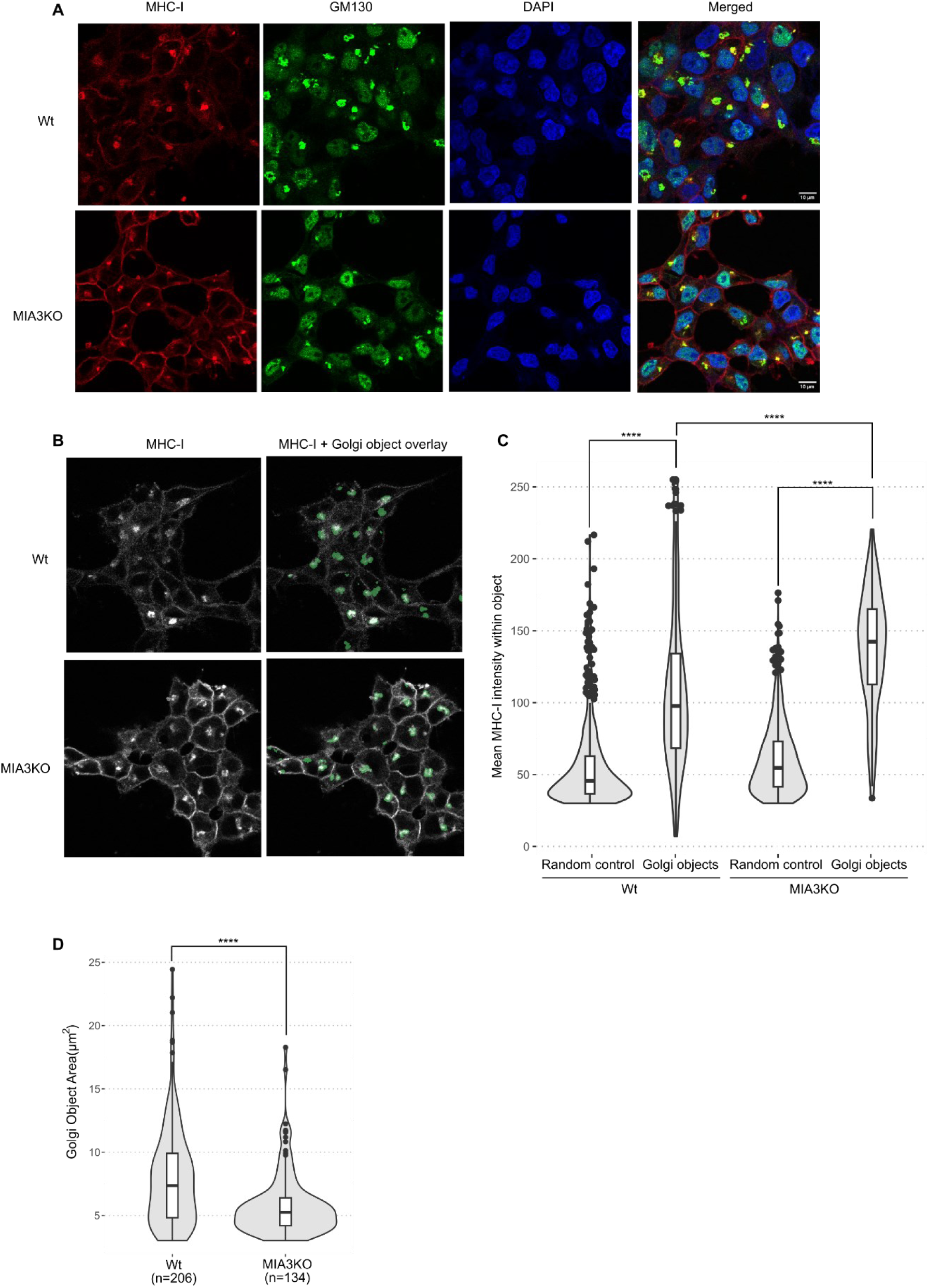
A KO of MIA3 in HEK293 cells reveals morphological changes in the Golgi but does not impact MHC-I presence within the organelle. (A) Confocal microscopy of WT and MIA3KO cells fixed and stained with DAPI, anti-MHC-I and anti-GM130 antibodies. Golgi are visible as bright green structures. (B) Golgi objects (in green) were isolated from images using Fiji and Weka trainable segmentation and overlaid on their corresponding MHC-I signal. MHC-I bright spots correspond with green Golgi objects in both cell types. (C) MHC-I bright spots correspond to GM130-positive objects. Colocalization was assessed in Fiji by measuring the intensity of the MHC-I signal within previously segmented Golgi objects. Random objects within the cell were generated and measured as a control for each cell line. A Mann-Whitney test (p<0.0001) was performed between groups to assess statistical significance. (D) Golgi object areas were measured using Fiji. 206 objects were identified in WT cells and 134 in MIA3KO cells. Violin plots show the distribution of Golgi object size. A Mann-Whitney test (p<0.001) revealed smaller Golgi in MIA3KO cells.

### HLA-A2-TurboID biotinylates target proteins more efficiently than HLA-A2-BioID2 and reveals different targets

The above-described results highlight the usefulness of our experimental approach for characterizing the MHC-I proxiome. However, the absence of proteins such as Tapasin and TAP1/2, which are part of the peptide-loading complex, suggests that there are limitations to the technique. Two parameters that can be easily modified are the nature of the biotin ligase and the length of the linker separating it from the MHC molecule. Thus, following the same principle as previously, the TurboID enzyme was fused to the intracytoplasmic domain of HLA-A2. To allow for more freedom of motion, a longer linker was added, this time consisting of four repeats of the

GGGGS motif. (Figure 8A) This new fusion protein was then expressed in HEK293 cells. The presence of surface MHC-I and intracellular biotin were measured by flow cytometry after a biotin exposition ranging from 15 to 60 minutes and compared to BioID2 construct in conditions used above. We find that biotinylation is increased in HLA-A2-TurboID-transfected cells when compared to HLA-A2-BioID2-transfected cells with as little as 15 minutes of exposure to biotin (Figure 8B). These observations are in line with previous literature comparing the two enzymes. ^14,15^ Biotinylated proteins from HLA-A2-TurboID transfected cells exposed to biotin for 60 minutes were collected as previously described for BioID2 and analysed using MS. Proteomic data was filtered using the same method as above, including the removal of HLA-DMα-BioID2 hits. The same procedure was performed for an associated triplicate of HLA-A2-BioID2 samples without retaining only hits present in 7 or more samples overall to allow for direct comparison between biotin ligases. Of the 946 proteins retained in the triplicate BioID2 and TurboID triplicate samples, 96 were exclusively found in BioID2 samples, 418 only in TurboID and 454 were found with both enzymes. (Figure 8C). Of the 454 common proteins, detection levels differed between the enzymes. Figure 8D shows proteins found in significantly different proportions as black-bordered circles in a volcano plot comparing expression for both enzymes. All 454 proteins are detailed in table S4. To further characterize targets from figure 8, three lists of proteins were input to the PANTHER algorithm ^27,28^ to detect enrichment for gene ontology terms along the 3 main categories. The first category containing proteins found only or significantly more (p<0.05) in BioID2 samples. The second containing those found only or significantly more in TurboID samples and the third, those labelled in equal proportions by both enzymes. Resulting fold-enrichment numbers were compared between the proteins more present in BioID2 vs. in TurboID samples. To avoid bias towards GO terms with a small number of members, only terms with 80 or more genes were considered. Figure 9 presents GO terms linked to pathways and locations of interest for MHC-I and reveals differences in detection between the two biotin ligases. For example, on one hand, BioID2 favored the identification of proteins linked to glycosylation and compartments such as the autophagosomes, lysosomes, and lipid droplets. On the other hand, the TurboID configuration skewed targets towards vesicular transport with greater enrichment for proteins found in the trans-Golgi network and endosomes, as well as transport vesicles, such as those coated with COPII, which are known to transport MHC-Is. ^42^

**Figure 8:**
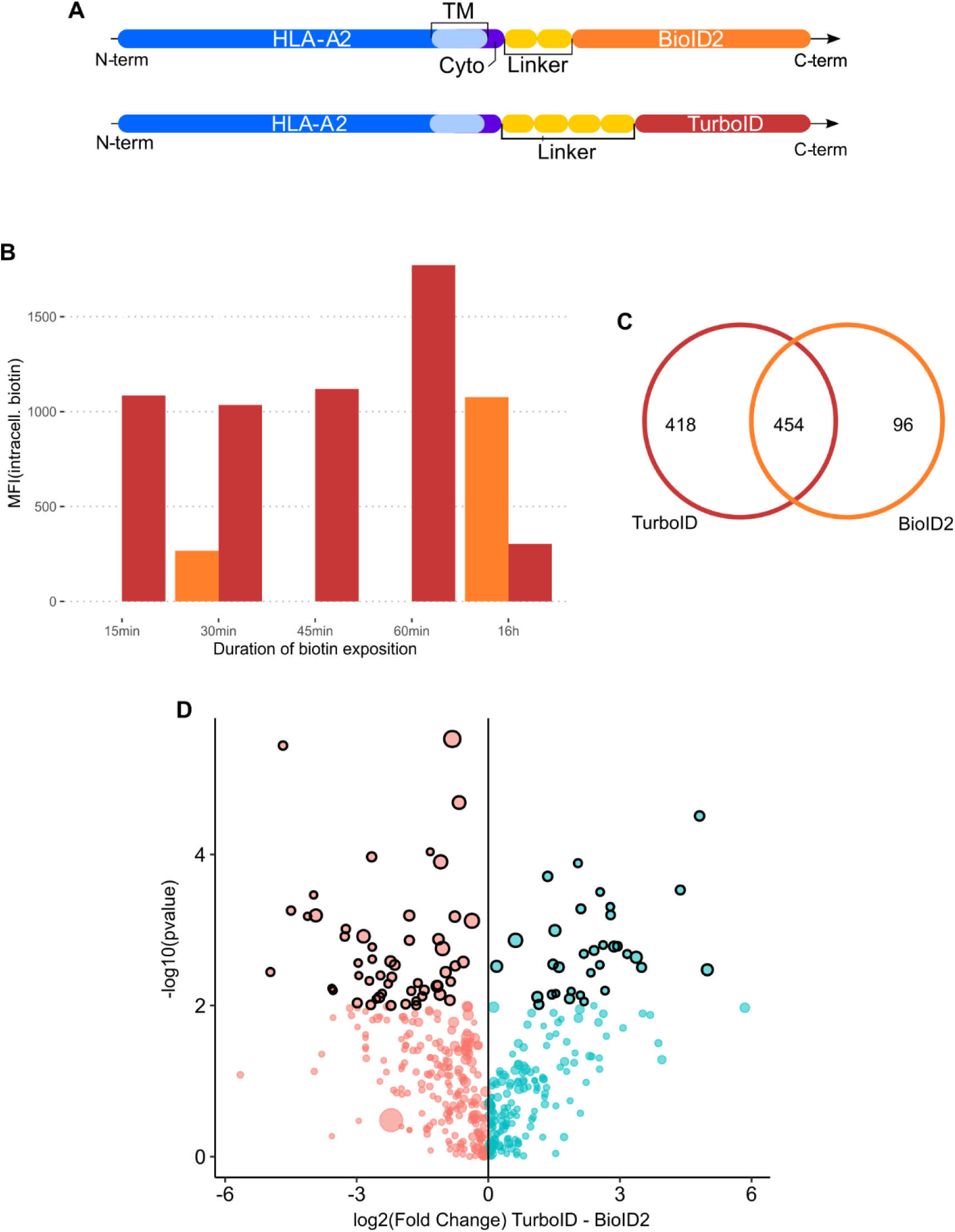
The HLA-A2-TurboID construct generates more and different biotinylated targets compared to HLA-A2-BioID2. **(A) Schematic representation of the HLA-A2-TurboID construct and comparison to HLA-A2 BioID2.** The HLA-A2 coding DNA sequence was fused at its C-terminus to the TurboID enzyme, originating from *E.coli*. The HLA-A2 termination codon was changed for a flexible ser-gly linker composed of 4 repeats of a GGGGS motif. **(B) Detection of intracellular biotin produced by HLA-A2-TurboID at different biotin exposure durations and comparison to HLA-A2-BioID2.** HEK293 cells were transfected with either HLA-A2-TurboID or HLA-A2-BioID2 as previously described. Cells that received TurboID (in red) were exposed to 70uM biotin for durations between 15 and 60 minutes, while BioID2 transfected cells (in orange) were exposed to biotin for 30 minutes or 16 hours. Expression of intracellular biotin and surface MHC-I was measured by flow cytometry. A ratio of the MFI values for intracellular biotin and surface MHC-I was calculated to assess the biotinylation level independent of transfection efficiency. **(C) Distribution of mass spectrometry hits between labeling enzymes.** Venn diagram organizing the 946 MS hits by their presence in BioID2 and TurboID triplicate samples. **(D) Differences in protein quantities found in both BioID2 and TurboID.** Volcano plot organizing proteins found in both BioID2 and TurboID samples by log2(fold change) on the X axis and −log10(p-value) on the Y axis, both for the comparison of TurboID and BioID2. Red circles represent hits found more abundantly in BioID2 samples, while blue circles represent hits found more abundantly in TurboID samples. Circle size represents average protein expression over all samples. Circles with a black border have a significantly (p<0.005) different expression according to a T-test with Welch’s correction.

**Figure 9:**
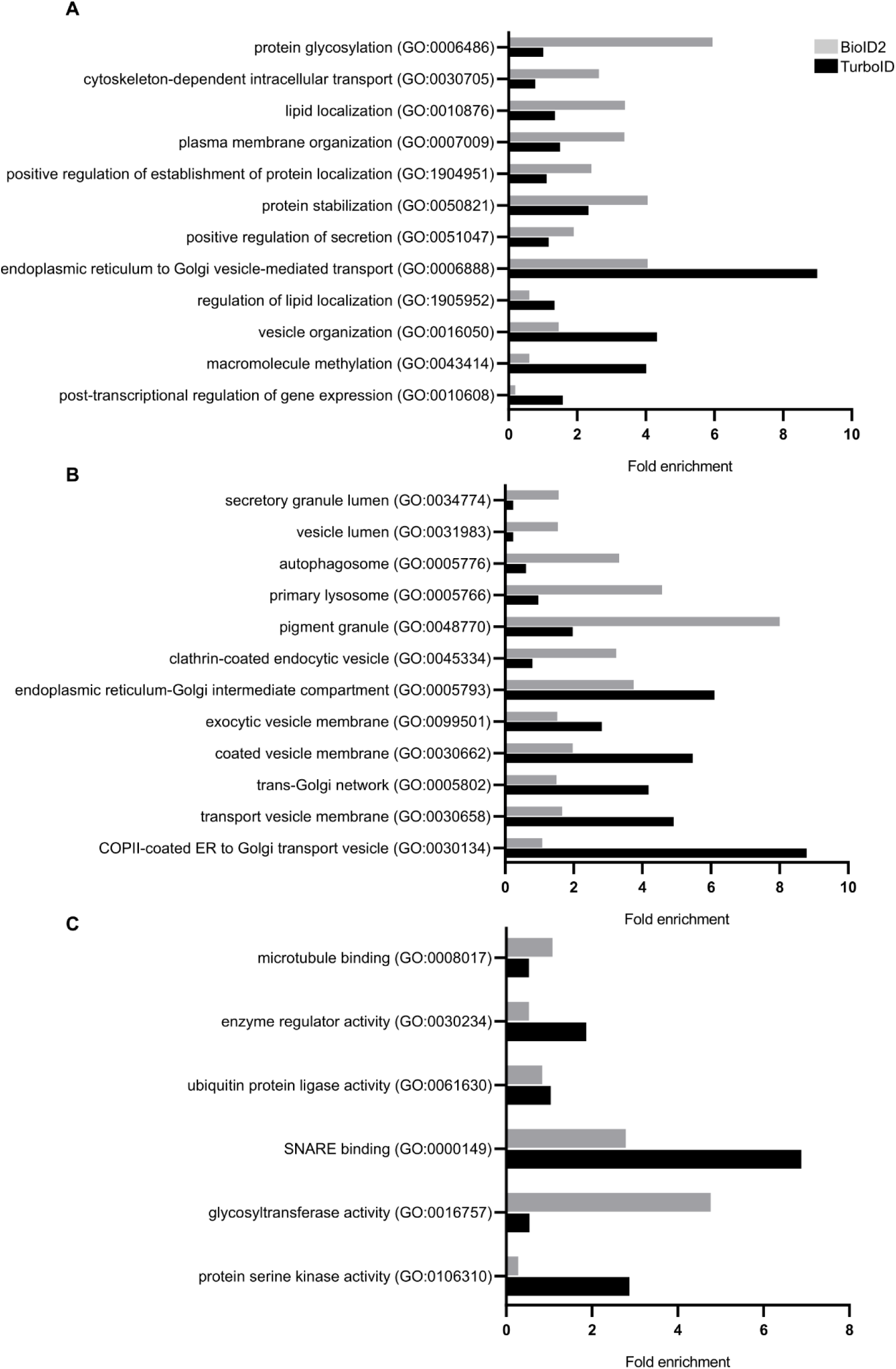
HLA-A2-BioID2 and HLA-A2-TurboID reach different targets according to gene ontology analysis. Proteins described in figure 7 were separated into 3 lists: those found only or significantly more (P<0.05) in BioID2 samples, those found only or significantly more in TurboID samples and those found in equal proportions by both enzymes. Of all GO terms, those related to MHC-I synthesis, transport, or function with significant differences between enzymes were plotted according to fold enrichment in each protein list when compared to the human genome. A plot was generated for each of the 3 main GO categories with: (A) Biological process, (B) Cellular component and (C) Molecular function. Only GO terms with more than 80 members were considered.

### TurboID allowed for the identification of MHC-I interactions implicated in downstream signaling after crosslinking

MHC-I crosslinking is known to activate various cell type-specific signaling pathways leading to a multitude of effects, ranging from apoptosis in T cells to cytokine production in macrophages and epithelial cells. ^43^ However, many of these downstream pathways and the implicated proteins are not well characterized. To validate our TurboID approach to the study of MHC reverse signaling, MHC-Is on HLA-A2-TurboID-transfected HEK293 cells were crosslinked with a combination of non-saturating concentrations of the W6-32 primary antibody and a GAM secondary antibody. After 1h at 37 degrees in biotin-supplemented medium, cells were lysed, and biotinylated material was processed for mass spectrometry. A total of 2974 individual hits were discovered in a triplicate experiment, which were narrowed down to 840 using the filtration methods described previously for both BioID2 and TurboID. (Supplementary Table 5, Figure 10A) Of these 840 proteins, 91 were significantly differently biotinylated when comparing mock- and W6/32-crosslinked samples. (Figure 10A,B and Table S5) Gene ontology (GO) enrichment analysis was then performed for these proteins and compared to the rest of the dataset. Fold enrichment for all GO biological process terms in both crosslinked and mock-crosslinked samples were plotted against each other to reveal terms overrepresented in one subset over another. Beyond this, Figure 11A highlights GO terms that are children of 3 parent terms: Transport, Metabolic Processes and Signaling to show that GO terms more enriched in crosslinked samples tend to relate to metabolism and transport, and not to signaling. Figure 11B compares the prevalence of specific GO terms as a percentage of the total genes for each condition.

**Figure 10:**
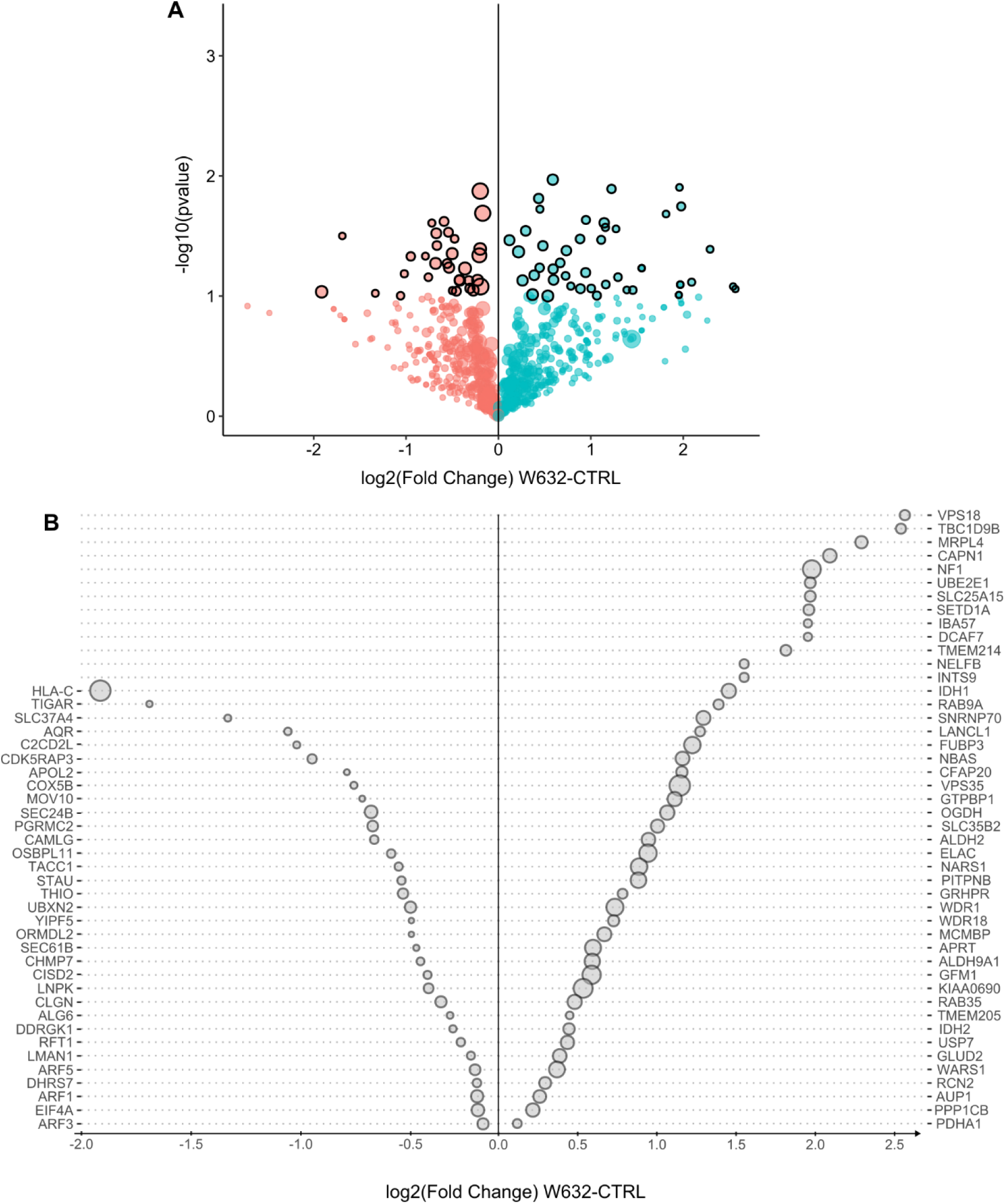
HLA-A2-TurboID reveals proteins potentially implicated in MHC-I reverse signaling. **(A)** Volcano plot showing the distribution of 824 hits found in both W6/32 and mock crosslinked samples organized by log2(Fold Change) on the X axis and −log10(pvalue) on the Y axis. 2 one-tailed Welch-corrected t-tests (p<0.05) were performed for each protein to identify targets significantly more and less biotinylated in W6/32 crosslinked samples, which are represented by black outlines. Average biotinylation across all samples is represented by the size of the circles. **(B)** The 91 targets with significantly different biotinylation were plotted according to log2(Fold Change) with their associated gene symbol. Genes on the right Y axis are found more in W6/32 crosslinked samples while genes on the left Y axis are found less in W6/32-crosslinked samples when compared to the isotype control-crosslinked samples. Once again, average biotinylation levels are represented by circle size.

**Figure 11:**
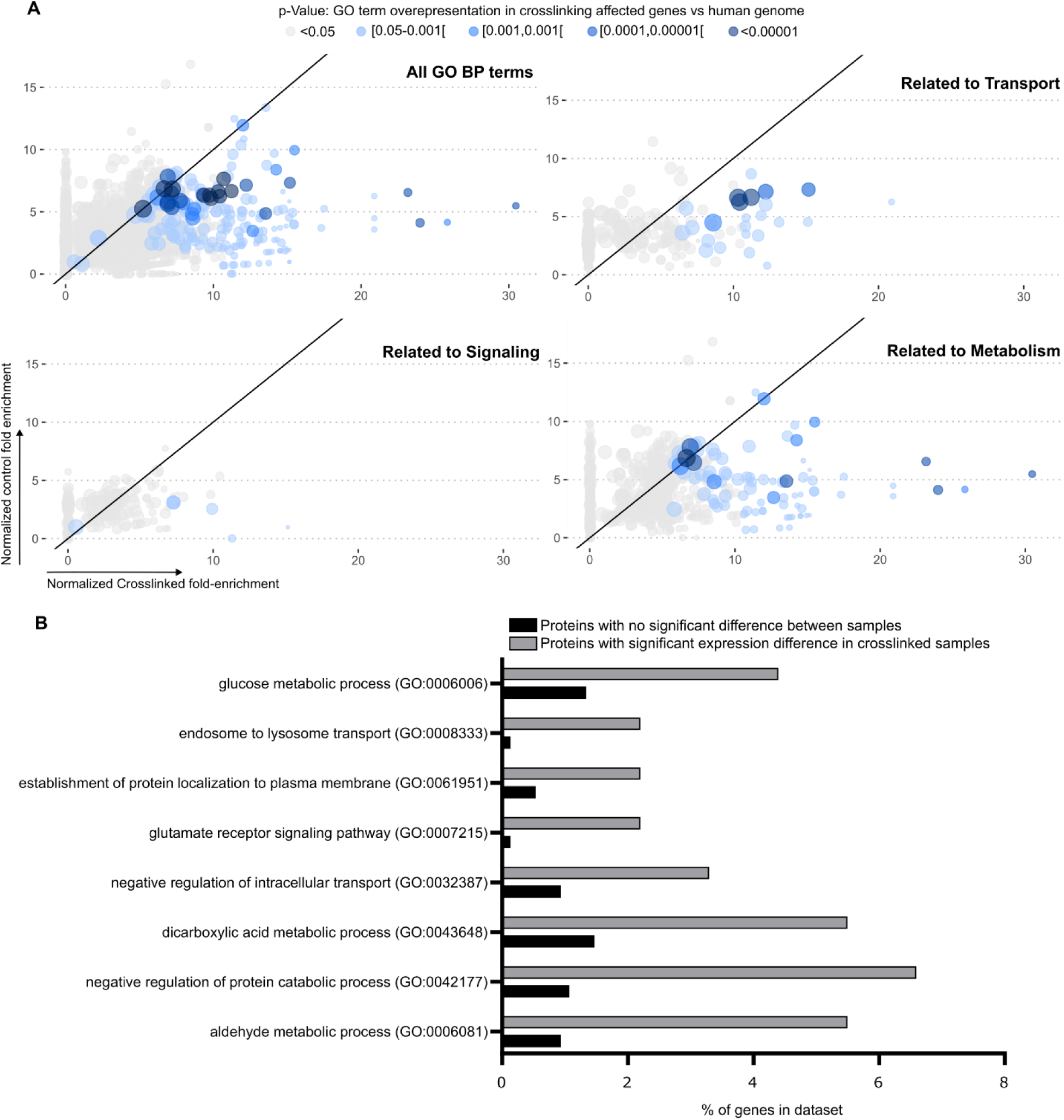
Proteins enriched with HLA-A2 cross-linking are mostly related to cellular transport and metabolic processes. (A) Gene ontology analysis of all biological process terms with between 80 and 2000 members. Terms are plotted on the X axis according to their fold enrichment in crosslinked samples and on the Y axis according to their fold enrichment in control samples both in regard to the human genome. Colors show statistical significance of enrichment in crosslinked samples. Top left plot shows all biological process terms. Top right shows only those that fall under Transport (GO:0006810), bottom left only those that fall under either Signalling (GO:0023052) or Regulation of Signalling (GO:0023051) and bottom right shows those that fall under Metabolic Processes (GO:0008152). Point size corresponds to the number of entries in each GO term. The black diagonal line represents equal fold-enrichment in both conditions. (B) Relevant GO terms with wide differences in representation in crosslinked and control samples are plotted according to the percentage of genes within each condition represented by each GO term.

Remarkably, despite all the changes in the proxiome triggered by W6/32, a bulk RNA-seq analysis conducted 24h after crosslinking showed a single differentially expressed gene, which codes for a proline-specific tRNA. (Figure S3) It is improbable that major transcriptional differences occurred at earlier time points and that cells had returned to normal after 24h. Most likely, in HEK293 cells, crosslinked MHC-Is induced proximal signaling events and shifts that trigger cell-autonomous, non-transcriptional metabolic regulation, amongst other pathways.

## DISCUSSION

Antigen presentation to T cells is a complex, highly regulated process. For more than four decades, the molecular basis behind the capture and display of self or non-self antigens has been the subject of intense research ^44,45^. That proteins must be digested into peptides to form a complex with MHC molecules and that the T cell receptor interaction is MHC restricted were two key discoveries in our understanding of T cell activation. ^46,47^ Over the years, the dissection of the MHC class I and II antigen presentation pathways highlighted important particularities in the acquisition of peptides. The fundamental difference between the loading of endogenous versus exogenous antigens resides in the subcellular location where peptide loading takes place, the former occurring within endocytic compartments often referred to as MIICs, while the latter occurs mostly within the endoplasmic reticulum. ^48^

While the general antigen presentation pathways are well-characterized, new players impacting directly or indirectly the trafficking and function of MHC molecules are discovered at a regular pace. Proximity ligation assays are powerful techniques used to uncover possible interactions between molecules in living cells. Our data extend the use of BioID to the study of the MHC-I’s cytoplasmic proxiome. Of the 4 main members of the PLC, only calreticulin was discovered in HLA-A2-BioID2 mass spectrometry data. However, it does not figure in the final 209 proteins, as it is eliminated due to its CRAPome score. While it was expected for ERp57 to be absent from our screen due to it residing on the lumenal side of the ER membrane, the absence of both TAP and Tapasin from our data merits further investigation as both are membrane proteins exposing a lysine residue in their intracytoplasmic domain, which could become biotinylated by BioID2. ^49^ It has been previously reported that, as opposed to TurboID, BioID is not efficient for the identification of ER residents. ^15,49^ Still, neither experimental approach fished out TAP and Tapasin in our hands using HEK293 cells. The absence of Tapasin could be due to the presence of the ligase, which occupies a 3×3.8nm space of the intracytoplasmic region where Tapasin could bind to MHC-I. ^13^ Interestingly, certain MHC-I alleles can be loaded and exported to the cell surface without the intervention of TAP or tapasin. HLA-A2 is not only capable of presenting Ags in a tapasin-independent manner, it was also reported as one of the alleles least affected by its absence. ^50^ Other cell types as well as other HLA class I alleles/isotypes will have to be tested before reaching definitive conclusions but our findings may highlight some important limitations of the proximity ligation approach. It would also be interesting to fuse the biotin ligase to the cytoplasmic region of TAP1/2 or Tapasin, or even to the N-terminal luminal end of MHC-1s. This last approach was successful in identifying some interactors for specific alleles of HLA-DP. ^51^

The use of HLA-DMα-BioID2 as a negative control has the effect of excluding many potential interactors located in the ER, as HLA-DMα is retained in this compartment without the presence of its complementary DMβ chain. (Table S1) The non-physiological accumulation of unfolded DMα chains most likely results in the biotinylation of random ER proteins. Without taking the control HLA-DMα-BioID2 into account, many known HLA-A2 interactors are revealed. These include TMX1, a thioredoxin which is known to associate with the MHC-I heavy chain and promotes peptide quality control. ^7^ Also, BAP31 associates with the peptide loading complex and is known to influence MHC-I transport. ^6,9,52^ Other examples include ESYT1 and VAPA, which, together with BAP31, have been shown to co-precipitate with MHC-I in dendritic cells. ^9^

The stringent filtering strategy described in figure 4A aimed at narrowing down the proxiome principally to targets of mature HLA-A2 and to reflect the post-ER proxiome of MHC-1s. Still, many proteins of the short list are ER residents and may play a role in the early steps of MHC-I folding and sorting. We were particularly intrigued by the presence of MIA3 in 10 out of 12 HLA-A2 proxiome experiments. This protein is known to modulate secretory vesicle export in the context of collagen, which is mediated by COPII-coated vesicles. ^37,39^ More specifically, MIA3 forms ring-like structures around ER-Golgi vesicles, which are required for the formation of COPII rings around the budding vesicles. ^37–39^ MHC-I also transits through COPII vesicles and HLA-A2 has previously been shown to be loaded into these vesicles by the Sec23/24 complex following an interaction of Sec23 with MHC-I’s C-terminal valine residue. ^53^ Similarly, an interaction between Bap31 and MHC-I has been associated with loading into ERGIC vesicles, which include COPII compartments, as well as with MHC-I quality control. ^6,52,54^ Interestingly, all 3 of these proteins have been found consistently in our HLA-A2-BioID2 experiments. MIA3 has also previously been shown to bind Sec23 and other members of the vesicle budding complex, such as Sec16 and NRZ, via different domains and to act as a filter for membrane lipids entering the newly formed vesicle. ^55,56^. We could envision a model where MIA3 loss leads to less stringent MHC-I quality control, resulting in more MHC-I export to the Golgi and the cell surface. Changes in Golgi morphology, which is expected in MIA3KO cells as smaller Golgi are consistent with the previously described ERGIC disruption caused by such a knockout. ^57^ Interestingly, our confocal microscopy data revealed that the presence of MHC-I is increased within these smaller Golgi, which is consistent with a quality control dysregulation hypothesis.

While BioID2 is an efficient tool to identify protein-protein interactions, it is somewhat limited by its long incubation time, which makes it difficult to study events that occur in a shorter timespan.^15^ Firstly, it was expected that TurboID would yield a greater number of unique interactors than BioID2 as its increased biotinylation efficiency was showed in previous studies. ^15^ Figure 7C shows a 1.58-fold increase in unique identified proteins between a triplicate HLA-A2-TurboID experiment and an identical HLA-A2-BioID2 triplicate processed simultaneously. However, it is difficult to attribute this improvement in detection to the TurboID enzyme alone, as the presence of a longer linker in this fusion protein should theoretically allow for a greater range of motion and thus more biotinylated targets. Indeed, a recent study based on APEX2 revealed drastically different pictures in subsequent experiments across labeling timepoints ranging from a few seconds to a few hours. ^58^ We therefore cannot hope to capture the entirety of the HLA-A2 proxiome with a single BioID or TurboID experiment. In line with these findings, the function and cellular localization of the proteins identified preferentially by each biotin ligase seem to differ. Notably, gene ontology analysis of proteins found exclusively or significantly more in HLA-A2-BioID2 or TurboID2 samples reveals an overrepresentation of proteins associated with glycosylation in BioID2 samples (Figure7) with the appearance of enzymes such as MGAT1 and MGAT2, which are part of the classical pathway of protein N-glycosylation in mammals. ^59^ MHC-I heavy chains are glycosylated and the resulting glycan chains are cleaved by glucosidases to allow MHC-I interaction with calnexin and calreticulin, which explains the appearance of such proteins in our screen. ^59,60^ On the other hand, TurboID identified more targets associated with vesicular transport such as SNAREs and Rab GTPases (Figure 7). This difference between biotin-ligase targets could result from interaction time, as vesicle fusion is an extremely rapid process lasting some 200 microseconds. ^61^ The increased efficiency of TurboID could potentially enable the detection of such interactions, while BioID2 might be better suited to longer processes, such as the complex multi-step assembly of glycans.

Building on this rationale, we applied TurboID to uncover potential MHC-I partners implicated in reverse signalling following antibody crosslinking. The outcome is presumed to be highly dependent on cell type, leading to very different effects ranging from apoptosis to cytokine production and calcium release. ^43,62^. Among proteins found in our screen, SHP-2 (PTPN11), which was described as a downstream effector of MHC-I in epithelial cells and macrophages, was biotinylated equally in both crosslinked and mock-crosslinked conditions. ^63^ The Rho-A GTPase was also found at slightly higher, although not significantly different, levels in W6/32-crosslinked cells and is linked to MHC-I downstream signalling in endothelial cells which leads to cytoskeletal remodelling. ^64,65^ Beyond the known signaling partners, our analysis revealed 91 potential novel downstream partners of MHC-I. (Figure 9A,C) Among them, targets such as TIGAR, SLC37A4, and PDHA1 are related to various metabolic processes for which GO terms are highly enriched when comparing the 91 retained hits with the remainder of the data from this experiment. (Figure 9B,C) MHC molecules have previously been linked to glucose metabolism, with anti-MHC-I antibody treatment in melanoma cells leading to down-regulation of genes implicated in glycolysis. ^66^ While crosslinking does have a downstream effect on the MHC-I proxiome, it does not seem to lead to transcriptional regulation in our model. This could potentially be explained by the already poor capacity of HEK293 cells to metabolise glucose. ^67^

Our results highlight the need to combine complementary approaches to obtain a comprehensive view of the proxiome. Variables such as the choice of cell type, the feasibility of performing BioID in vivo in mice, the biotin ligase used, its orientation and linker length, and even the detergents employed all influence the outcome. By identifying MIA3 as a new regulator of MHC-I expression, we demonstrate the value of HLA-coupled biotin ligases for probing antigen presentation pathways in greater depth.

---

**FigureS1:**
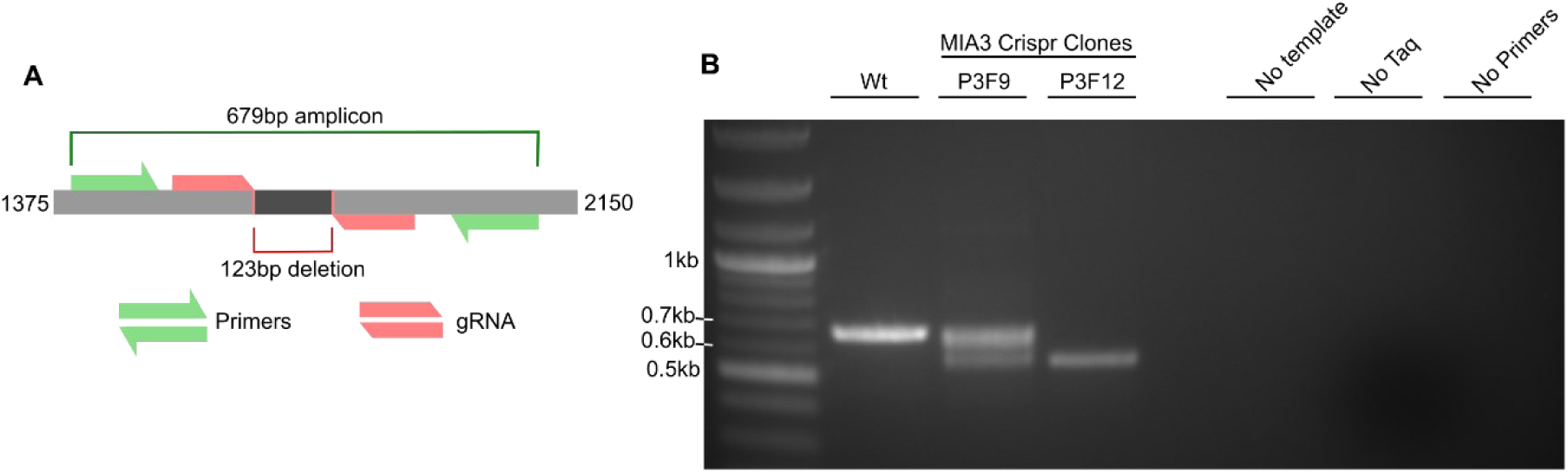
Confirmation of the MIA3 knockout by RT-PCR. (A) Schematic representation of the PCR method used to confirm MIA3 knockout in HEK293 cells. (B) PCR screening identified clone P3F12 as being MIA3KO. Bands of around 700bp indicate intact MIA3 genes, while bands of around 600bp indicate successful deletion in the MIA3 locus. Clone P3F9’s pattern indicates a deletion in only 1 of the 2 chromosomes.

**Figure S2:**
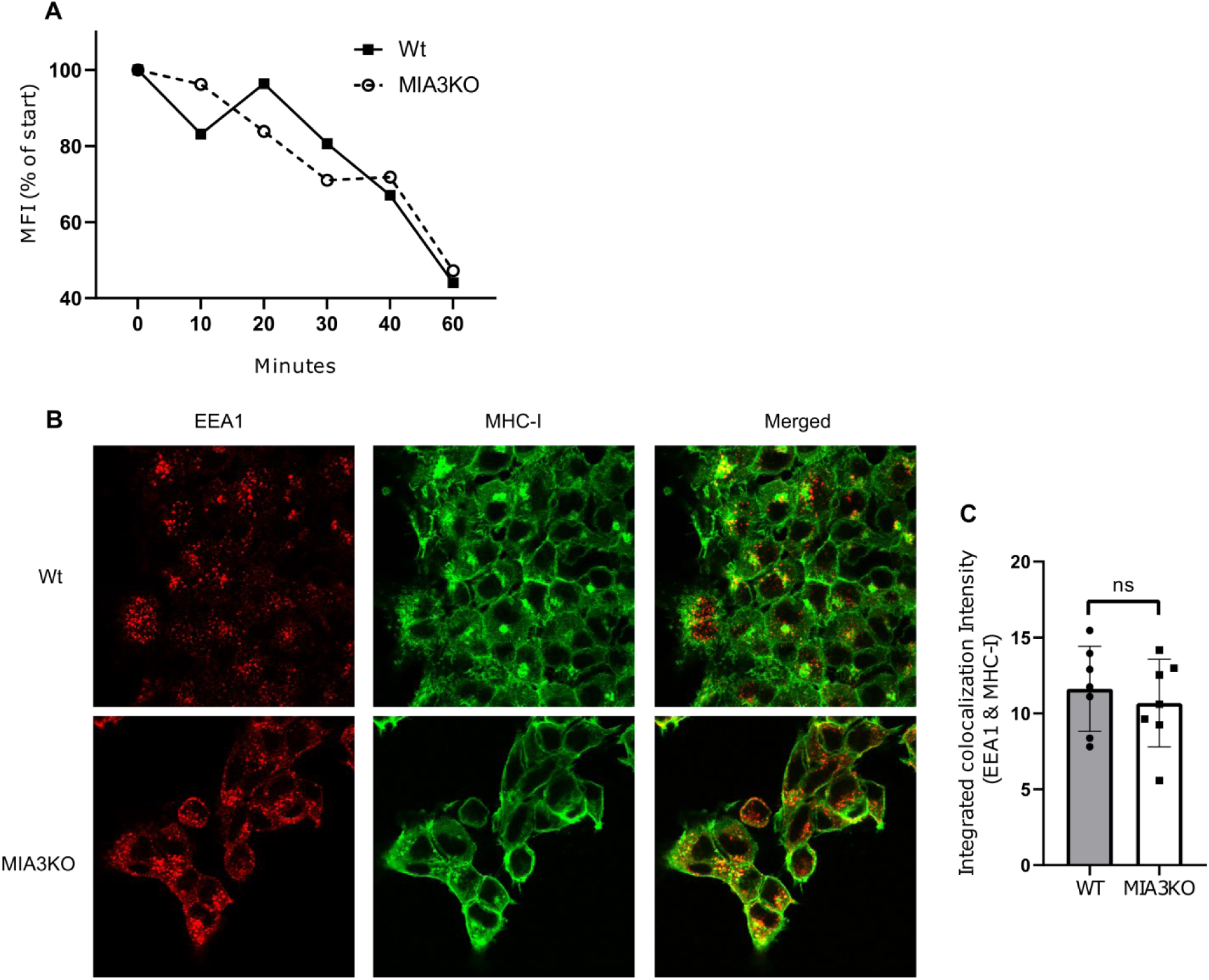
Endosomal localisation and internalisation of MHC-I are not affected by absence of MIA3. (A) Confocal microscopy staining of both the early endosomal marker EEA1 and MHC-I are similar in WT and MIA3KO cells. (B) MHC-I signal intensity within EEA1+ objects is not significantly different between WT and MIA3KO cells. (C) Flow cytometry measures of W6/32 internalization show no real difference between WT and MIA3KO cells.

**Figure S3:**
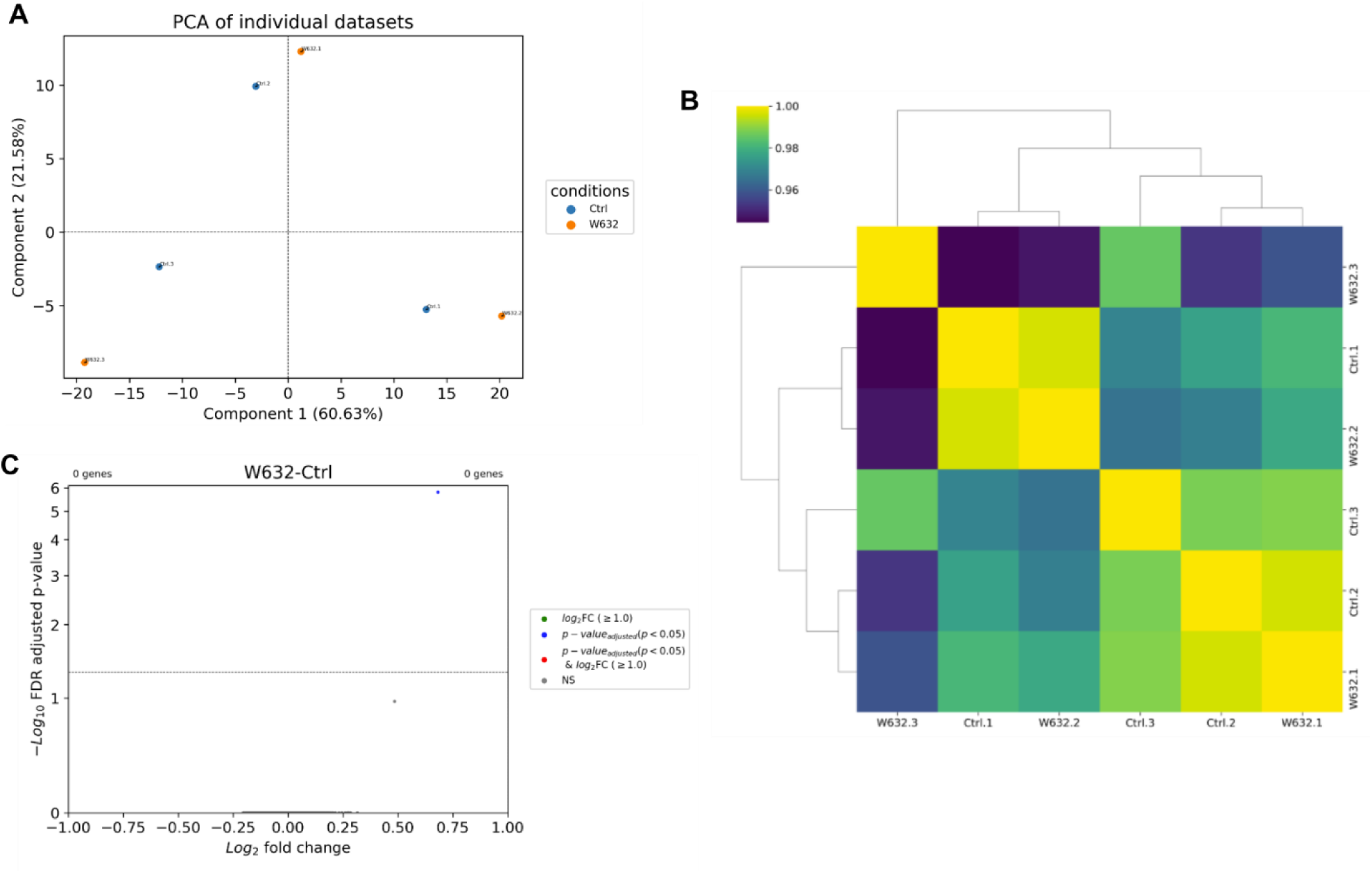
Transcriptome is not affected by W6/32 crosslinking after 24h. (A) PCA analysis of RNA-seq results for HEK293 cells crosslinked with W6/32 or Isotype control antibody show no difference between conditions (B) Sample correlation heatmap shows no distinction between conditions (C) Volcano plot showing a single differentially expressed gene between crosslinked and non-crosslinked cells.

## ACKNOWLEDGEMENTS

Special thanks to Dr. Nicole Bernard for kindly providing the K-562 cells, Dr. Michel Desjardins for providing various antibodies, Dr. Joseph Sodroski for providing cells and antisera, Dr. Nicolas Stifani for his expertise in microscopy and Dr. Armelle Le Campion for her help with flow cytometry. We would also like to thank the IRCM’s molecular biology and bioinformatics platforms for their help regarding transcriptomic data. J.T. holds the *Saputo* research chair.

## Authorship Contributions

W. M. designed experiments, performed experiments, discussed results, and wrote the paper; E.L. performed microscopy experiments and image analysis; D.F. performed the mass spectrometry analysis and wrote the associated protocol; S. Z. provided key reagents; J.T. designed experiments, discussed results, and wrote the paper. All authors approved the final version of the manuscript.

## References

1 Blees, A. et al. Structure of the human MHC-I peptide-loading complex. Nature 551, 525–528 (2017). 10.1038/nature24627

2 Fisette, O., Schröder, G. F. & Schäfer, L. V. Atomistic structure and dynamics of the human MHC-I peptide-loading complex. Proceedings of the National Academy of Sciences 117, 20597–20606 (2020). 10.1073/pnas.2004445117

3 Montealegre, S. & van Endert, P. M. Endocytic Recycling of MHC Class I Molecules in Non-professional Antigen Presenting and Dendritic Cells. Frontiers in Immunology 9 (2019).

4 Adiko, A. C., Babdor, J., Gutiérrez-Martínez, E., Guermonprez, P. & Saveanu, L. Intracellular Transport Routes for MHC I and Their Relevance for Antigen Cross-Presentation. Frontiers in Immunology 6, 335 (2015). 10.3389/fimmu.2015.00335

5 Paulsson, K. M. et al. Association of tapasin and COPI provides a mechanism for the retrograde transport of major histocompatibility complex (MHC) class I molecules from the Golgi complex to the endoplasmic reticulum. The Journal of Biological Chemistry 277, 18266–18271 (2002). 10.1074/jbc.M201388200

6 Abe, F., Van Prooyen, N., Ladasky, J. J. & Edidin, M. Interaction of Bap31 and MHC Class I Molecules and Their Traffic Out of the Endoplasmic Reticulum 1. The Journal of Immunology 182, 4776–4783 (2009). 10.4049/jimmunol.0800242

7 Matsuo, Y., Masutani, H., Son, A., Kizaka-Kondoh, S. & Yodoi, J. Physical and functional interaction of transmembrane thioredoxin-related protein with major histocompatibility complex class I heavy chain: redox-based protein quality control and its potential relevance to immune responses. MBoC 20, 4552–4562 (2009). 10.1091/mbc.e09-05-0439

8 Cebrian, I. et al. Sec22b regulates phagosomal maturation and antigen crosspresentation by dendritic cells. Cell 147, 1355–1368 (2011). 10.1016/j.cell.2011.11.021

9 Barends, M. et al. Dynamic interactome of the MHC I peptide loading complex in human dendritic cells. Proceedings of the National Academy of Sciences 120, e2219790120 (2023). 10.1073/pnas.2219790120

10 Roux, K. J., Kim, D. I., Burke, B. & May, D. G. BioID: A Screen for Protein-Protein Interactions. Curr Protoc Protein Sci 91, 19 23 11–19 23 15 (2018). 10.1002/cpps.51

11 Roux, K. J., Kim, D. I., Raida, M. & Burke, B. A promiscuous biotin ligase fusion protein identifies proximal and interacting proteins in mammalian cells. J. Cell Biol 196, 801–810 (2012). https://doi.org/jcb.201112098 [pii];10.1083/jcb.201112098 [doi]

12 Liu, X., Salokas, K., Weldatsadik, R. G., Gawriyski, L. & Varjosalo, M. Combined proximity labeling and affinity purification-mass spectrometry workflow for mapping and visualizing protein interaction networks. Nat. Protoc 15, 3182–3211 (2020). 10.1038/s41596-020-0365-x

13 Kim, D. I. et al. An improved smaller biotin ligase for BioID proximity labeling. Mol Biol Cell 27, 1188–1196 (2016). 10.1091/mbc.E15-12-0844

14 Branon, T. C. et al. Efficient proximity labeling in living cells and organisms with TurboID. Nat Biotechnol 36, 880–887 (2018). 10.1038/nbt.4201

15 May, D. G., Scott, K. L., Campos, A. R. & Roux, K. J. Comparative Application of BioID and TurboID for Protein-Proximity Biotinylation. Cells 9, 1070 (2020). 10.3390/cells9051070

16 Barnstable, C. J. et al. Production of monoclonal antibodies to group A erythrocytes, HLA and other human cell surface antigens-new tools for genetic analysis. Cell 14, 9–20 (1978). https://doi.org/0092-8674(78)90296-9 [pii]

17 Kelly, A. P., Monaco, J. J., Cho, S. & Trowsdale, J. A new human HLA class II-related locus, DM. Nature 353, 571–573 (1991).

18 Burr, M. L. et al. An Evolutionarily Conserved Function of Polycomb Silences the MHC Class I Antigen Presentation Pathway and Enables Immune Evasion in Cancer. Cancer Cell 36, 385–401.e388 (2019). 10.1016/j.ccell.2019.08.008

19 Balthazard, R. et al. A PROXIMITY LIGATION SCREEN IDENTIFIES SNAT2 AS A NOVEL TARGET OF THE MARCH1 E3 UBIQUITIN LIGASE. bioRxiv, 2024.2009.2016.613264 (2024). 10.1101/2024.09.16.613264

20 Mellacheruvu, D. et al. The CRAPome: a contaminant repository for affinity purification–mass spectrometry data. Nat Methods 10, 730–736 (2013). 10.1038/nmeth.2557

21 Shannon, P. et al. Cytoscape: a software environment for integrated models of biomolecular interaction networks. Genome Research 13, 2498–2504 (2003). 10.1101/gr.1239303

22 Doncheva, N. T. et al. Cytoscape stringApp 2.0: Analysis and Visualization of Heterogeneous Biological Networks. J Proteome Res 22, 637–646 (2023). 10.1021/acs.jproteome.2c00651

23 Szklarczyk, D. et al. The STRING database in 2023: protein-protein association networks and functional enrichment analyses for any sequenced genome of interest. Nucleic Acids Res 51, D638–D646 (2023). 10.1093/nar/gkac1000

24 Li, M., Li, D., Tang, Y., Wu, F. & Wang, J. CytoCluster: A Cytoscape Plugin for Cluster Analysis and Visualization of Biological Networks. Int J Mol Sci 18, 1880 (2017). 10.3390/ijms18091880

25 Ge, S. X., Jung, D. & Yao, R. ShinyGO: a graphical gene-set enrichment tool for animals and plants. Bioinformatics 36, 2628–2629 (2020). 10.1093/bioinformatics/btz931

26 Dhanda, S. K. et al. IEDB-AR: immune epitope database-analysis resource in 2019. Nucleic Acids Research 47, W502–W506 (2019). 10.1093/nar/gkz452

27 Thomas, P. D. et al. PANTHER: Making genome-scale phylogenetics accessible to all. Protein Sci 31, 8–22 (2022). 10.1002/pro.4218

28 Mi, H. et al. Protocol Update for large-scale genome and gene function analysis with the PANTHER classification system (v.14.0). Nat Protoc 14, 703–721 (2019). 10.1038/s41596-019-0128-8

29 Stirling, D. R. et al. CellProfiler 4: improvements in speed, utility and usability. BMC Bioinformatics 22, 433 (2021). 10.1186/s12859-021-04344-9

30. Schindelin, J., et al. Fiji: an open-source platform for biological-image analysis. Nat Methods 9, 676-682 (2012). 10.1038/nmeth.2019

31 Arganda-Carreras, I. et al. Trainable Weka Segmentation: a machine learning tool for microscopy pixel classification. Bioinformatics 33, 2424–2426 (2017). 10.1093/bioinformatics/btx180

32 Chen, X., Zaro, J. L. & Shen, W.-C. Fusion protein linkers: property, design and functionality. Adv Drug Deliv Rev 65, 1357–1369 (2013). 10.1016/j.addr.2012.09.039

33 Krshnan, L., van de Weijer, M. L. & Carvalho, P. Endoplasmic Reticulum-Associated Protein Degradation. Cold Spring Harb Perspect Biol 14 (2022). 10.1101/cshperspect.a041247

34 Wälchli, S. et al. Invariant chain as a vehicle to load antigenic peptides on human MHC class I for cytotoxic T-cell activation. Eur J Immunol 44, 774–784 (2014). 10.1002/eji.201343671

35. Oughtred, R., et al. The BioGRID database: A comprehensive biomedical resource of curated protein, genetic, and chemical interactions. Protein Science : A Publication of the Protein Society 30, 187-200 (2021). 10.1002/pro.3978

36 Matko, J., Bushkin, Y., Wei, T. & Edidin, M. Clustering of class I HLA molecules on the surfaces of activated and transformed human cells. Journal of Immunology (Baltimore, Md.: 1950) 152, 3353–3360 (1994).

37 Raote, I. et al. TANGO1 assembles into rings around COPII coats at ER exit sites. The Journal of Cell Biology 216, 901–909 (2017). 10.1083/jcb.201608080

38 Raote, I., Saxena, S., Campelo, F. & Malhotra, V. TANGO1 marshals the early secretory pathway for cargo export. Biochimica et Biophysica Acta (BBA) - Biomembranes 1863, 183700 (2021). 10.1016/j.bbamem.2021.183700

39 Reynolds, H. M., Zhang, L., Tran, D. T. & Ten Hagen, K. G. Tango1 coordinates the formation of endoplasmic reticulum/Golgi docking sites to mediate secretory granule formation. The Journal of Biological Chemistry 294, 19498–19510 (2019). 10.1074/jbc.RA119.011063

40 Lamb, J. E., Ray, F., Ward, J. H., Kushner, J. P. & Kaplan, J. Internalization and subcellular localization of transferrin and transferrin receptors in HeLa cells. J Biol Chem 258, 8751–8758 (1983).

41 Tortorella, S. & Karagiannis, T. C. Transferrin Receptor-Mediated Endocytosis: A Useful Target for Cancer Therapy. The Journal of Membrane Biology 247, 291–307 (2014). 10.1007/s00232-014-9637-0

42 Spiliotis, E. T., Manley, H., Osorio, M., Zuniga, M. C. & Edidin, M. Selective Export of MHC Class I Molecules from the ER after Their Dissociation from TAP. Immunity 13, 841–851 (2000).

43 Muntjewerff, E. M., Meesters, L. D., Bogaart, G. v. d. & Revelo, N. H. Reverse Signaling by MHC-I Molecules in Immune and Non-Immune Cell Types. Frontiers in Immunology 11 (2020).

44 Pamer, E. & Cresswell, P. Mechanisms of MHC class I--restricted antigen processing. Annu. Rev. Immunol 16, 323–358 (1998).

45 Brunnberg, J. et al. Dual role of the peptide-loading complex as proofreader and limiter of MHC-I presentation. Proc Natl Acad Sci U S A 121, e2321600121 (2024). 10.1073/pnas.2321600121

46 Zinkernagel, R. M. & Doherty, P. C. Restriction of in vitro T cell-mediated cytotoxicity in lymphocytic choriomeningitis within a syngeneic or semiallogeneic system. Nature 248, 701–702 (1974). 10.1038/248701a0

47 Buus, S. et al. Interaction between a “processed” ovalbumin peptide and Ia molecules. Proceedings of the National Academy of Sciences of the United States of America 83, 3968–3971 (1986). 10.1073/pnas.83.11.3968

48 Blum, J. S., Wearsch, P. A. & Cresswell, P. Pathways of antigen processing. Annu. Rev. Immunol 31, 443–473 (2013). 10.1146/annurev-immunol-032712-095910 [doi]

49 Afshar, N., Black, B. E. & Paschal, B. M. Retrotranslocation of the Chaperone Calreticulin from the Endoplasmic Reticulum Lumen to the Cytosol. Mol Cell Biol 25, 8844–8853 (2005). 10.1128/MCB.25.20.8844-8853.2005

50 Lewis, J. W., Sewell, A., Price, D. & Elliott, T. HLA-A*0201 presents TAP-dependent peptide epitopes to cytotoxic T lymphocytes in the absence of tapasin. European Journal of Immunology 28, 3214–3220 (1998). 10.1002/(SICI)1521-4141(199810)28:10<3214::AID-IMMU3214>3.0.CO;2-C

51 Anczurowski, M. et al. Chaperones of the class I peptide-loading complex facilitate the constitutive presentation of endogenous antigens on HLA-DP84GGPM87. Journal of Autoimmunity 102, 114–125 (2019). 10.1016/j.jaut.2019.04.023

52 Paquet, M.-E., Cohen-Doyle, M., Shore, G. C. & Williams, D. B. Bap29/31 influences the intracellular traffic of MHC class I molecules. Journal of Immunology (Baltimore, Md.: 1950) 172, 7548–7555 (2004). 10.4049/jimmunol.172.12.7548

53 Cho, S., Ryoo, J., Jun, Y. & Ahn, K. Receptor-Mediated ER Export of Human MHC Class I Molecules Is Regulated by the C-Terminal Single Amino Acid. Traffic 12, 42–55 (2011). 10.1111/j.1600-0854.2010.01132.x

54 Ladasky, J. J. et al. Bap31 Enhances the Endoplasmic Reticulum Export and Quality Control of Human Class I MHC Molecules1. The Journal of Immunology 177, 6172–6181 (2006). 10.4049/jimmunol.177.9.6172

55 Raote, I. et al. TANGO1 membrane helices create a lipid diffusion barrier at curved membranes. eLife 9, e57822 (2020). 10.7554/eLife.57822

56 Aridor, M. A tango for coats and membranes: New insights into ER-to-Golgi traffic. Cell Reports 38, 110258 (2022). 10.1016/j.celrep.2021.110258

57 McCaughey, J. et al. A general role for TANGO1, encoded by MIA3, in secretory pathway organization and function. Journal of Cell Science 134, jcs259075 (2021). 10.1242/jcs.259075

58 Susa, K. J. et al. A spatiotemporal map of co-receptor signaling networks underlying B cell activation. Cell Rep 43, 114332 (2024). 10.1016/j.celrep.2024.114332

59 Ryan, S. O. & Cobb, B. A. Roles for major histocompatibility complex glycosylation in immune function. Semin Immunopathol 34, 10.1007/s00281-00012-00309-00289 (2012). https://doi.org/10.1007/s00281-012-0309-9

60 Trombetta, E. S. & Helenius, A. Lectins as chaperones in glycoprotein folding. Current Opinion in Structural Biology 8, 587–592 (1998). 10.1016/S0959-440X(98)80148-6

61 Haluska, C. K. et al. Time scales of membrane fusion revealed by direct imaging of vesicle fusion with high temporal resolution. Proceedings of the National Academy of Sciences 103, 15841–15846 (2006). 10.1073/pnas.0602766103

62 Eyford, B. A. et al. Outside-in signaling through the major histocompatibility complex class-I cytoplasmic tail modulates glutamate receptor expression in neurons. Sci Rep 13, 13079 (2023). 10.1038/s41598-023-38663-z

63 Xu, S. et al. Constitutive MHC class I molecules negatively regulate TLR-triggered inflammatory responses via the Fps-SHP-2 pathway. Nat Immunol 13, 551–559 (2012). 10.1038/ni.2283

64 Lepin, E. J., Jin, Y. P., Barwe, S. P., Rozengurt, E. & Reed, E. F. HLA class I signal transduction is dependent on Rho GTPase and ROK. Biochem Biophys Res Commun 323, 213–217 (2004). 10.1016/j.bbrc.2004.08.082

65 Ziegler, M. E., Souda, P., Jin, Y. P., Whitelegge, J. P. & Reed, E. F. Characterization of the endothelial cell cytoskeleton following HLA class I ligation. PloS one 7, e29472 (2012). 10.1371/journal.pone.0029472

66 Peppicelli, S. et al. Potential Role of HLA Class I Antigens in the Glycolytic Metabolism and Motility of Melanoma Cells. Cancers (Basel) 11 (2019). 10.3390/cancers11091249

67 Abaandou, L., Quan, D. & Shiloach, J. Affecting HEK293 Cell Growth and Production Performance by Modifying the Expression of Specific Genes. Cells 10 (2021). 10.3390/cells10071667

